# Dissolution-Controlled Nanocrystalline Rifapentine Formulation for Tuberculosis Treatment

**DOI:** 10.64898/2026.08.16.745059

**Authors:** Nisha S. Barge, Yeswanth C. Kalapala, Pranshu Rajurkar, Ameya A. Dravid, Naveen K. Bhukya, Rituparna Saha, V Sanjay, Harinath Chakrapani, Rachit Agarwal

## Abstract

Current tuberculosis (TB) treatment suffers from drawbacks such as long regimens, high pill burden and side effects leading to non-adherence and poor treatment outcomes. Dissolution-controlled drug depot formulation with high drug loading is a clinically successful drug delivery strategy. Such depots reduce the dosing frequency for treatments requiring daily administration, thereby improving treatment adherence and compliance. However, dissolution-controlled depots for first-line TB drugs have not been demonstrated due to their high solubility and high dose requirements. In this study, we overcame this challenge by developing injectable, extended-release, dissolution-controlled depots of nanocrystalline rifapentine (NCRPT), microcrystalline rifapentine (MCRPT) and amorphous rifapentine microparticles (ARPT) with more than 75% loading. Crystalline formulations resulted in much slower depot dissolution compared to amorphous formulations. A single intramuscular (IM) injection of NCRPT in mice resulted in therapeutic serum concentrations for over a week. We then demonstrated the efficacy of NCRPT in both pre-exposure prophylaxis and therapeutic models of mice TB. NCRPT administered at 60 mg/kg once every two weeks demonstrated excellent efficacy in a mouse model of TB infection. In each case, a ∼ 4-log-fold reduction in lung bacterial load compared to untreated mice was observed. These results open new avenues for developing LAI formulations of TB drugs and could improve patient compliance and TB management.

---

Tuberculosis (TB) is a major global challenge with 10.7 million reported cases in 2024 and resulted in 1.23 million deaths [1]. The WHO-approved chemotherapy for TB includes several drugs, with rifamycins being the prominent antibiotics for first-line treatment, administered orally for 4-6 months. Despite being effective, the high mortality burden indicates certain drawbacks of the current treatment regimens and hinders the complete eradication of TB. The most significant drawback is the long treatment duration. The inherent ability of *Mycobacterium tuberculosis* (Mtb), the causative agent of TB, to enter a non-replicating state *in vivo* consequently requires a high pill burden and longer duration of antibiotic therapy for complete eradication [2]. The use of antibiotics for a longer duration causes side effects such as fatigue, nausea, etc., leading to reduced quality of life, prompting non-adherence [3]. Non-adherence to TB treatment results in poor treatment outcomes and the emergence of drug resistance, aggravating the challenge [4]. Studies carried out in developing countries observe non-adherence as the most prominent reason hindering successful treatment, highlighting the need for treatments with shorter durations and less frequent dosing [5]. TB preventive therapy (TPT) is prescribed for individuals at a high risk of developing active TB disease, which includes people with HIV and household contacts of TB patients. WHO recommends several TPT regimens ranging from 1-9 months [6]. Although highly successful in preventing TB infections in 90% of the cases [7], TPT also suffers from non-adherence due to longer duration and frequent dosing [8,9]. These drawbacks of the current treatment highlight the urgent need for shorter treatment regimens, reduced dosage frequency and fewer side effects, ensuring treatment adherence.

Advances in drug delivery strategies could hold a potential solution to the challenges posed in TB prevention and treatment. Developing delivery systems with extended drug release could lead to reduced dosage frequency, improving treatment adherence. Several attempts have been made to achieve controlled and prolonged drug release using polymer-based formulations encapsulating isoniazid and rifampicin [10,11], delivering anti-TB drugs in alginate microspheres [12], nanoencapsulation of anti-TB drugs in polylactic-co-glycolic acid (PLGA) using spray drying technique [13], etc. Although these formulations extend the release of anti-TB drugs to some extent, the major limitation with these drug delivery systems is the low encapsulation efficiency and loading levels (ranging from 1-10 %). This necessitates an increased amount of polymers to be delivered to maintain therapeutic drug levels for longer durations *in vivo* [10,14]. Kim et al. developed an injectable *in-situ* forming polymer-drug implant for long-acting formulation of rifabutin with loading levels of ∼30-40% and releasing drug for up to 16 weeks, indicating the potential of advanced drug delivery strategies in reducing dosage frequency for TB treatment [15].

Developing long-acting injectable (LAI) formulations of drugs to achieve extended release is a clinically approved strategy used for improving patient compliance by reducing dosage frequency for the treatment of schizophrenia [16]. This strategy was also used in developing injectable nanocrystalline formulations of cabotegravir, allowing once every two months IM administration and was subsequently approved for pre-exposure prophylactic therapy for HIV [17,18]. Despite being a very promising drug delivery strategy, an anti-TB LAI formulation has been reported only for bedaquiline, a second-line drug [19], currently undergoing phase I trial for assessment of pharmacokinetics and safety in healthy individuals (EU Clinical Trial 2023-508810-41-00). Injectable formulations for first-line TB drugs have not been reported. Although several studies address the advantage of amorphousness over crystallinity for pharmaceutical drug formulations for increased bioavailability after oral administration [20, 21,22,23], the role of crystallinity/amorphousness in the case of dissolution-controlled long-acting injectable formulations in the context of TB has also not been reported. Many TB drugs are not suitable candidates for LAI formulations. Rifapentine (RPT), belonging to the rifamycin class of antibiotics, is a first-line anti-TB drug approved by the WHO for a shorter TB regimen in 2021 [24]. RPT, along with isoniazid, has also been recommended for a one-month TPT regimen [25]. RPT with low aqueous solubility, higher LogP, lower minimum inhibitory concentration (MIC) and longer pharmacokinetic half-life compared to other first-line TB drugs, forms the ideal candidate for developing LAI [26,27]. While LAI formulations of RPT have not been demonstrated, a previous study utilized dynamic oral dosing and estimated the required serum concentration (C_trough_) of RPT to be between 0.6 – 2 μg/mL for it to be bactericidal *in* vivo [28]. In this study, we synthesized and characterized various injectable long-acting rifapentine formulations: a nanocrystalline RPT formulation (NCRPT) and a microcrystalline RPT formulation (MCRPT) prepared via solvent-antisolvent precipitation and an amorphous RPT (ARPT) prepared via emulsion-based method. Owing to their higher free energy and lack of ordered structure, amorphous formulations are known to have faster dissolution compared to the crystalline formulations of the same drug [20,21,22,23], a fact utilized in improving the oral bioavailability of the drug. In our study, we demonstrate that the crystalline nature of the long-acting CRPT injectable formulation leads to a much longer-lasting drug depot compared to the ARPT in a mouse model. CRPT maintains RPT levels above 2 μg/ml in serum for over a week with a single IM administration. Once a week and once every two weeks, IM administration of CRPT has shown similar efficacy compared to daily orally administered RPT in pre-exposure prophylactic and therapeutic mouse models, indicating the translatability of the strategy in reducing dosage frequency and improving patient compliance.

## Results

### Synthesis and characterization of RPT formulations

We first developed methods to prepare high loading crystalline and amorphous formulations of rifapentine. To develop controlled-release, high-loading-level crystalline formulations of rifapentine, a solvent-antisolvent precipitation technique was used. A few solvent-antisolvent pairs were used along with varying amounts of RPT **(Table S1)**. The optimum solvent-antisolvent pair was selected based on crystal formation and high yield (∼50-60%). The crystals obtained were not injectable owing to their larger size (**Figure S1**). In order to reduce the size of crystals, RPT crystals were ultrasonicated (**Figure 1A**). The resulting nanocrystal RPT formulation (NCRPT), upon lyophilization, was used for physical characterization. Due to the non-spherical nature of crystalline particles, the size of RPT nanocrystals was quantified by Scanning Electron Microscope (SEM) imaging. The nanocrystals were observed to be 772.8 ± 356.9 nm in length and 602.01 ± 314.6 nm in width. To obtain an injectable microcrystal RPT formulation, RPT crystals were bath sonicated at a concentration of 10 mg/mL. The microcrystals were lyophilized and stored at room temperature (RT) for further use. This formulation was designated as microcrystal RPT formulation (MCRPT). The microcrystals of CRPT are observed to be 36.25 ± 18.11 µm in length and 24.6 ± 13.22 µm in width (**Figure 1B**). Amorphous rifapentine formulation (ARPT) was prepared by homogenization and resulted in spherical microparticles of an average diameter of 4.3 ± 1.83 µm (**Figure 1C**). Powder X-ray diffraction (pXRD) measurements revealed the crystalline nature of NCRPT and MCRPT, as indicated by the sharp peaks in the pXRD spectrum (**Figure 1D and 1E**). These sharp peaks are absent from the spectrum for ARPT, indicating the amorphousness of the formulation (**Figure 1F**). The crystalline character of NCRPT and MCRPT is further confirmed by the differential scanning calorimetry (DSC) measurements. A sharp melting peak around 160 °C (melting temperature, T_m_) demonstrates crystallinity in the NCRPT formulation (**Figure 1G**) and MCRPT formulation (**Figure 1H)**. The melting peak is absent from the DSC thermograph of ARPT (**Figure 1I**).

**Figure 1.**
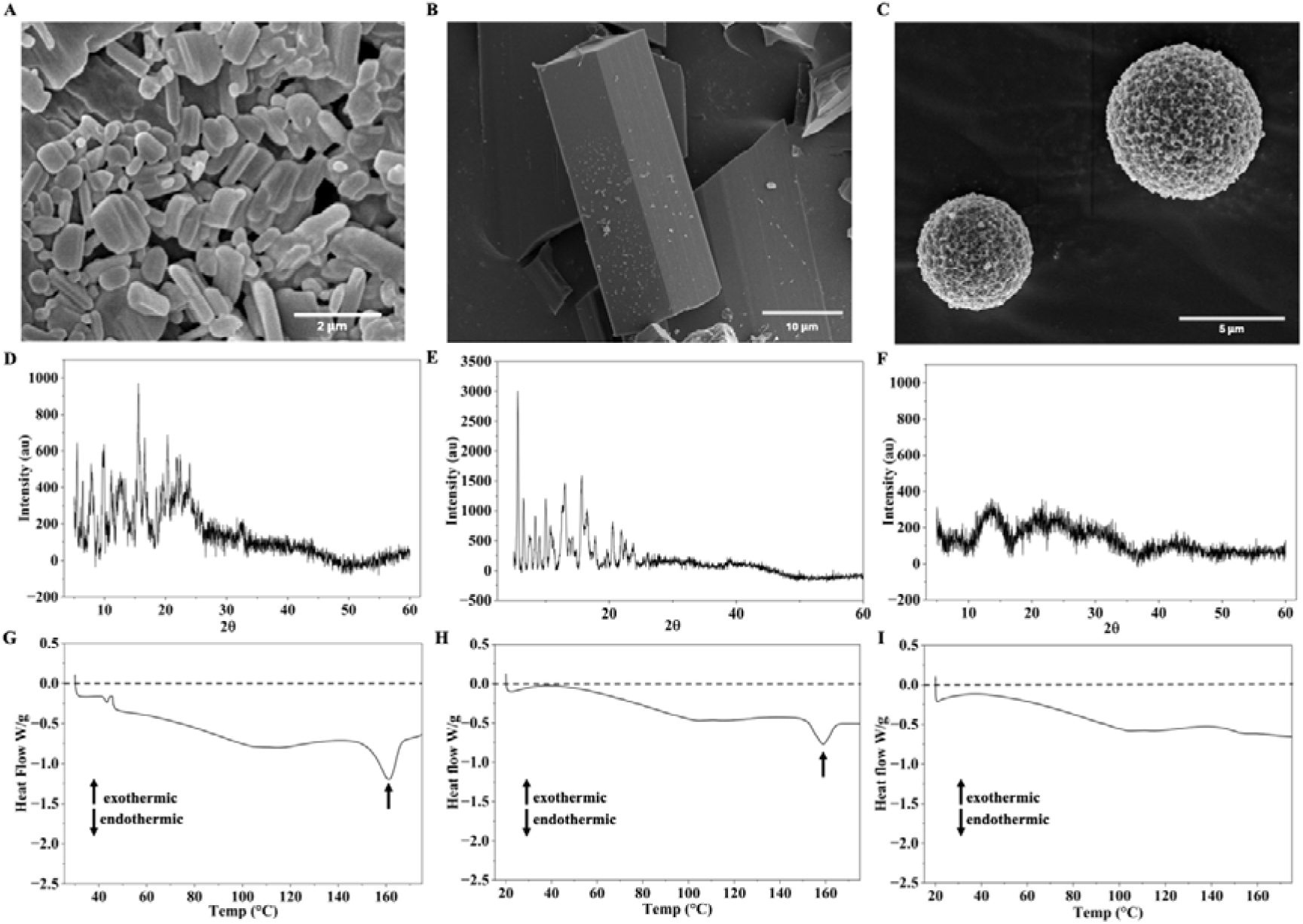
Characterization of RPT formulations. (**A**) SEM image of RPT nanocrystals after ultrasonication (scale bar - 2 µm). (B) SEM image of RPT microcrystal formulation (scale bar – 10 µm). (**C**) SEM image of ARPT formulation (scale bar – 5 µm). **(D**) pXRD analysis of NCRPT, indicating the crystallinity of RPT nanocrystals. **(E)** pXRD analysis of MCRPT, indicating the crystallinity of RPT microcrystals. **(F)** pXRD analysis of ARPT, indicating the amorphousness of the formulation. **(G)** DSC thermogram of NCRPT, black arrow indicates the melting peak. (**H**) DSC thermogram of MCRPT, black arrow indicates the melting peak. **(I)** DSC thermogram of ARPT.

RPT loading levels in NCRPT and MCRPT formulations was quantified by dissolving the formulation in methanol and determining the RPT content through absorbance. RPT loading levels were observed to be 87.7 ± 1.42 % in NCRPT and 94.05 ± 1.67 % in MCRPT, which allows a large amount of drug to be administered in low-volume injections. To our knowledge, this is the highest reported loading level of RPT in any nano/micro-formulation. RPT loading levels in ARPT were quantified by dissolving the formulation in DMSO and determining the RPT content through absorbance. RPT loading levels were observed to be 76.2 ± 17.5 %.

### *In vitro* dissolution study with NCRPT, MCRPT and ARPT

To determine the effect of crystallization and size of crystals on the dissolution of RPT from our formulations, an *in vitro* dissolution study was performed. Dialysis cups (10 kDa membrane cut-off) with NCRPT, MCRPT and ARPT formulations at a concentration of 1 mg/mL in 1X PBS were suspended in a 1X PBS sink (sink changed every 24 h). Samples from dialysis cups were analyzed by dissolving in methanol to quantify RPT amount every 24 h. Faster dissolution was observed in the case of ARPT compared to both NCRPT and MCRPT (**Figure 2A**). This indicates the potential of crystalline formulations in reducing dissolution, leading to the possibility of forming longer-lasting depots *in vivo*. Despite higher surface-to-volume ratio in NCRPT compared to MCRPT, we do not see a significant difference in dissolution rates for the two formulations, suggesting higher contribution of crystallinity in controlling dissolution rates compared to particle size.

**Figure 2.**
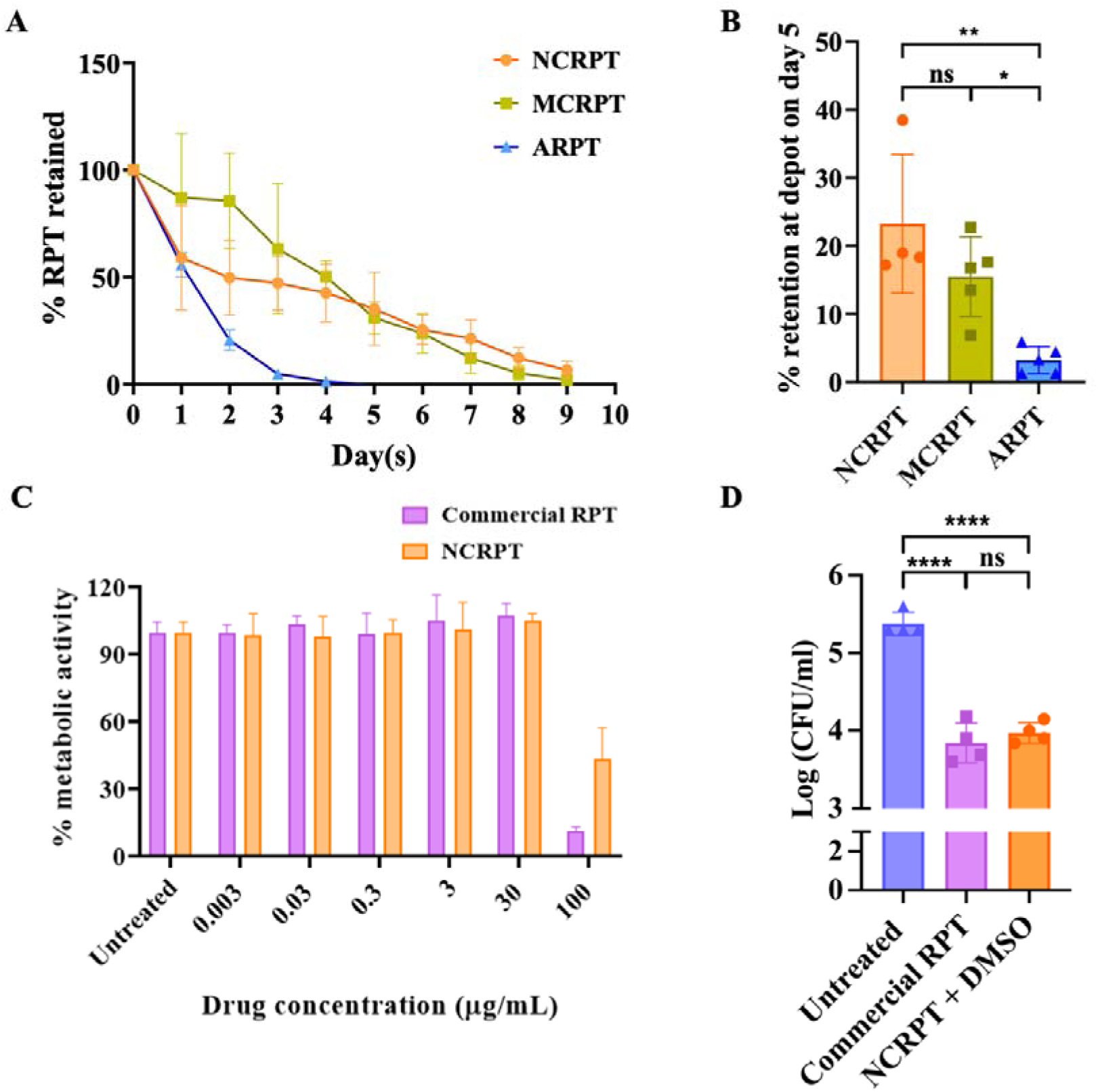
Crystalline formulation forms a longer-lasting drug depot in vivo compared to an amorphous formulation and retains its antibacterial efficacy. **(A)** In vitro assay to determine RPT released from NCRPT, MCRPT and ARPT (n = 3 distinct batches for all groups). **(B)** RPT amount at the site of injection on day 5 post-injection plotted as % retention (n=5 muscle tissues isolated from different animals for both groups). One data point for the CRPT group was excluded after outlier analysis using the Grubbs test. **(C)** WST-assay performed with THP-1 macrophages to determine the cytocompatibility of RPT nanocrystal formulation compared with the commercially available rifapentine as a control (n=4 distinct samples for both groups). **(D)** In vitro efficacy of RPT nanocrystal formulation tested in THP-1 macrophages infected with Mtb H37Rv (n=4 distinct samples for all groups). Data in the graph represents mean ± SD. p-value < 0.05 was considered significant. For A, p-value = 0.0056 was determined by an unpaired t-test with Welch’s correction. For **B**, p-values were determined by ordinary one-way ANOVA followed by post-hoc Tukey’s multiple comparison tests. p-value < 0.05 was considered significant. ****p < 0.0001 for control vs commercial RPT (5 μg/mL), ****p < 0.0001 for control vs CRPT + DMSO (5 μg/mL) and ns = non-significant.

### *In vivo* depot dissolution study with NCRPT, MCRPT and ARPT

To determine the controlled and sustained dissolution of our RPT formulations in an animal model, we administered NCRPT, MCRPT and ARPT formulations to mice in the caudal thigh (rifapentine dosage of 60 mg/kg). RPT present in the depot was extracted using methanol and quantified at day 0 and day 5 post-injection from the isolated muscle. The weight chart recorded for animals shows no toxicity for any of the formulations (**Figure S2)**. Similar to our in vitro studies, at day 5, RPT retention was comparable between NCRPT (23.2 ± 10.2 %) and MCRPT (15.5 ± 5.83), while it was significantly lower for ARPT (3.25 ± 2.01%) (**Figure 2B)**. Despite having a higher surface-to-volume ratio in NCRPT nanocrystals than in micron-sized particles in ARPT, these results clearly showed that crystalline formulations significantly slow down the dissolution compared to amorphous formulations *in vivo*. Although different surfactants were required to be used for different techniques to prepare crystalline and amorphous formulations, the high loading levels of rifapentine suggest that surfactants are unlikely to be a major contributor to the differences observed in dissolution rates. In addition, surfactants are only expected to be on the surface of the crystals, which are likely to be solubilized as the surface of particles dissolves. No significant difference was observed in the day 5 depot concentrations for NCRPT and MCRPT, as was seen in the *in vitro* study. It was also observed that RPT microcrystals in the MCRPT formulation tend to settle quickly upon resuspension during injections, leading to non-uniformity during animal studies. As a result, we have focused on NCRPT formulation for all subsequent assays.

### Injectability and stability of NCRPT formulation

To assess the injectability of 60 mg/kg NCRPT through a 27 G × ½” needle, the RPT concentration in the NCRPT formulation before and after injection was determined. No significant difference was observed in the RPT concentration before and after injection (**Figure S3**). To assess any changes in the formulation characteristics after injection, the size of the nanocrystals was determined before and after injection using SEM. As reported earlier, the size of nanocrystals before injection was observed to be 772.8 ± 356.9 nm in length and 602.01 ± 314.6 nm in width (n=300 from three distinct batches). After injection, the size was observed to be 662.31 ± 270.74 nm in length and 495.22 ± 218.74 nm in width. These assays indicated the injectability and stability of NCRPT formulations. To assess the effect of crystallization on the dissolution of RPT, RPT crystals were incubated in 1X PBS at a concentration of 0.01 mg/ml in a 96-well plate and individual crystals were monitored by microscopy over 6 hours. Dissolution of two representative crystals is shown in **Figure S4**. It was observed that the area of the crystals reduces over time, while still maintaining the shape of the crystals, indicating surface dissolution. The assay was performed on RPT crystals, as imaging the RPT nanocrystals over a period of 6 hours was not feasible with light microscopy. As seen in the case of these crystals, we expect similar surface dissolution of the drug from nanocrystals.

### *In vitro* cytotoxicity and efficacy of NCRPT formulation

To assess the cytotoxicity of RPT nanocrystals, the nanocrystals were incubated with THP-1 macrophages for 24 hours. Metabolic activity of the cells was determined using the WST reagent. RPT nanocrystals were well tolerated by macrophages, similar to the commercially available RPT (**Figure 2C**). Cytotoxicity was also assessed with human embryonic kidney (HEK-293) cells, and similar results were observed **(Figure S5)**.

The process of producing RPT nanocrystals exposes RPT to organic solvents and harsh treatments during homogenization and subsequent lyophilization. To assess whether the crystallization and formulation process affects the efficacy of RPT, we dissolved the NCRPT formulation in DMSO. THP-1 macrophages infected with a virulent strain of Mtb, H37Rv, were treated with dissolved RPT (diluted in PBS to the required concentration) for 48 hours. Infected THP-1 macrophages with no treatment formed the control, while treatment with commercially available RPT was used to compare the efficacy of RPT. Upon treatment with 5 µg/mL of drug for both treatment groups, a similar reduction in bacterial load was observed in both the treatment groups, with mean log_10_ CFU of 3.84 ± 0.26 and 3.97 ± 0.13 for commercial RPT and NCRPT (dissolved in DMSO), respectively, when compared with a mean log_10_ CFU of 5.375 ± 0.15 for the control. No significant difference was observed between treatment with commercially available RPT and DMSO dissolved RPT nanocrystal formulations, suggesting RPT was active against the bacteria after being exposed to various processes during formulation production (**Figure 2D**).

### Pharmacokinetics of NCRPT formulation

A pharmacokinetic study of RPT nanocrystal formulation was performed in male BALB/c mice. Animals were administered 60 mg/kg CRPT by IM injection in the caudal thigh muscle. The dose was determined considering the objective for long-term release of the drug. Doses beyond 60 mg/kg (corresponding to suspension beyond 15 mg/mL) started showing aggregation of particles. As a result, 60 mg/kg dose was used for all animal experiments. Post IM injection, serum samples were analyzed by LC-MS to assess RPT levels at different time intervals over 9 days. Weights of all the animals were recorded at regular intervals. The weight chart indicates no weight loss upon drug administration (**Figure 3A**). A previous study utilized dynamic oral dosing and estimated the required serum C_trough_ concentration of RPT to be between 0.6 – 2 μg/mL for it to be bactericidal *in vivo* [28]. Hence, we set 2 μg/mL as a target for our pharmacokinetic study. Remarkably, with just a single injection of our NCRPT formulation, RPT levels in serum remained considerably above 2 μg/mL until day 6, indicating sustained release of the drug for a week with a single administration (**Figure 3B**). The levels rose initially to 11 μg/mL by day 1, likely due to burst release, and then stabilized to 2-3 μg/mL for the rest of the week before dropping below the 2 μg/mL cut-off by day 9. Pharmacokinetic parameters for NCRPT were determined by using non-compartmental analysis (**Table S2**). Peak serum concentration (C_max_) was observed to be 11.15 μg/mL achieved in 24 hours post injection (T_max_). The calculated AUC (AUC_0–∞_) was 35.17 μg.day/mL. To determine RPT concentration in lungs, RPT levels in bronchoalveolar lavage fluid (BALF) were checked 24 hours post-dosing for animals dosed with either 20 mg/kg RPT orally (clinical dose for active TB) or 60 mg/kg NCRPT intramuscularly. No significant difference was observed in RPT levels in both groups, indicating NCRPT formulations were able to maintain RPT levels similar to oral dosage **(Figure S6)**.

**Figure 3.**
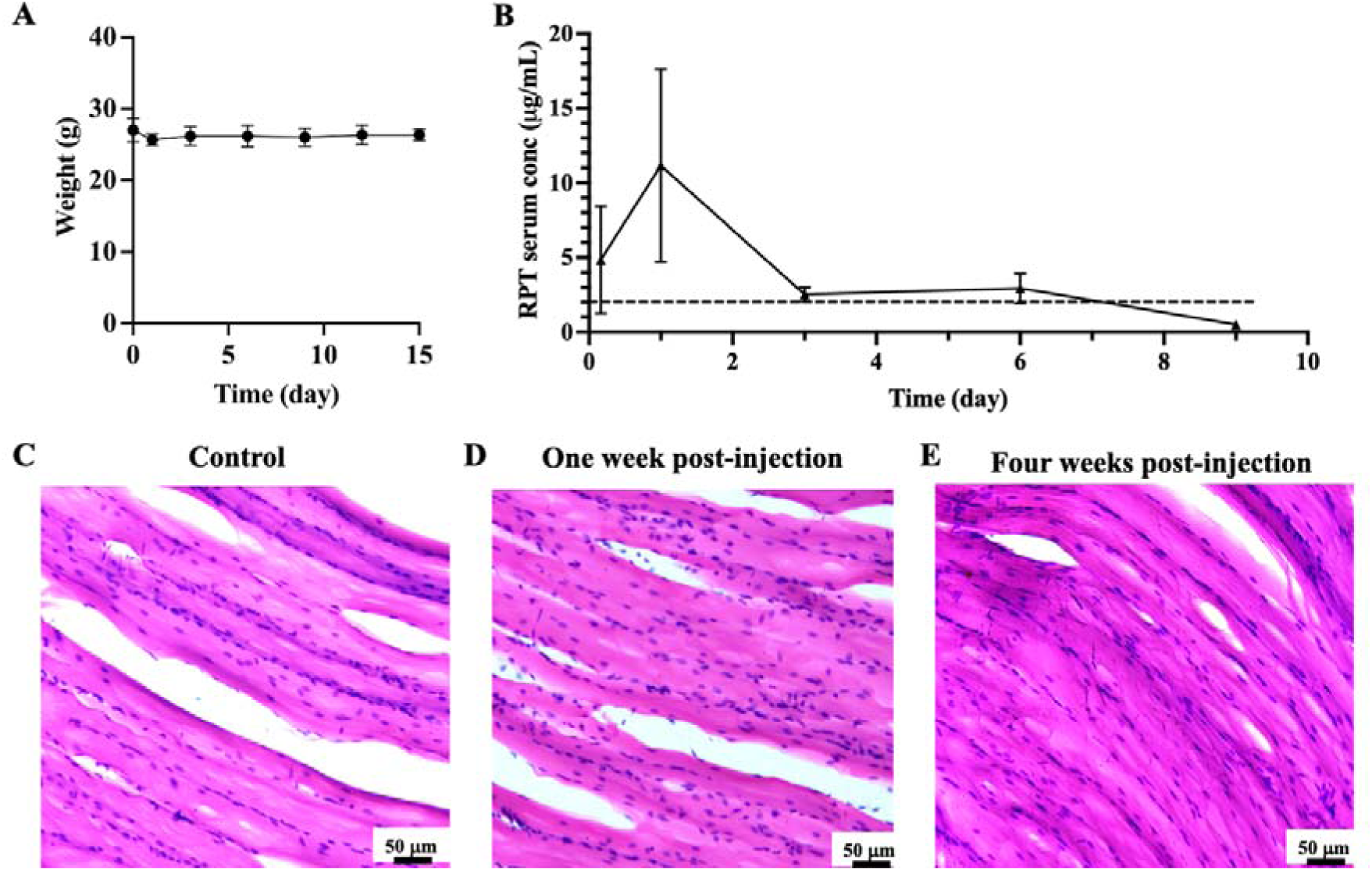
CRPT single IM administration releases the therapeutically required RPT level over one week and does not cause toxicity in mice. (**A**) Weight chart of mice. Weights were checked at regular intervals until day 15 after IM injection (n=4 different animals). (**B**) Pharmacokinetic analysis of RPT in mouse serum using LC-MS (n=4 serum samples from different animals). Dashed line at 2 μg/mL represents the reported therapeutically serum concentration of RPT. Data in the graph represent mean ± SD. Representative images of H and E stained muscle tissues. (**C**) Control. (**D**) One week and (**E**) Four weeks post-injection. Scale bar = 50 μm.

### NCRPT formulation safety analysis

Local toxicity due to IM injection of NCRPT formulation was assessed by doing histopathological analysis of the injected muscle with 60 mg/kg CRPT. Muscle tissues were cryo-sectioned, stained with hematoxylin and eosin (H&E) stain and visualized using a brightfield microscope. **Figure 3C-E** shows the representative stained muscle tissues one week (**Figure 3D**) and four weeks (**Figure 3E**) post-injection. Muscle tissue isolated from mice with no injection (**Figure 3C**) was used as a control. No significant immune infiltration was observed post-injection, indicating the safety of NCRPT formulations.

To evaluate for any systemic toxicity, blood serum samples isolated from untreated, single-dose orally administered RPT and single-dose intramuscularly injected NCRPT animals were tested for alanine aminotransferase (ALT), aspartate aminotransferase (AST) and total bilirubin levels. No significant difference was observed in the levels of ALT, AST and total bilirubin among untreated and orally administered RPT and intramuscularly injected NCRPT groups on either day 1 or day 6 post-drug administration (**Figure S7)**, indicating no systemic toxicity caused by NCRPT.

### Efficacy of NCRPT formulation in a pre-exposure prophylactic mouse model

RPT has been approved for a one-month oral regimen along with isoniazid for TB preventive therapy (TPT). WHO recommends TPT for people living with HIV and for household contacts of people with TB^5^. The *in vivo* efficacy of the NCRPT formulation was hence assessed in a pre-exposure prophylactic mouse model. Schematic of the experimental design and the different dose regimens included in the experiment are detailed in **Figure 4A**. The animals were either orally administered a daily dose (six out of seven days per week) of 20 mg/kg RPT (oral (6/7)) or NCRPT formulations injected intramuscularly (60 mg/kg) with a frequency of one injection per week (CRPT (1/7)) or one injection per two weeks (CRPT (1/14)). Three days after drug administration, animals were infected with Mtb H37Rv via aerosolization. Weights of the animals were recorded and showed no signs of toxicity of the formulations (**Figure 4B**). The day after infection, the mean log_10_ CFU was 3.34 ± 0.17. Upon four weeks of monitoring, all the treatment regimen groups showed a reduced lung bacterial load of 1.47 ± 0.38, 1.06 ± 0.15 and 1.03 ± 0.08 for oral (6/7), NCRPT (1/7) and NCRPT (1/14) groups respectively, when compared with the control group (infected, untreated) that harbored a bacterial load of 6.6312 ± 0.33 log_10_ CFU **(Figure 4C)**. Animals administered with RPT orally at a frequency of 6 days per week harboured 1.092 ± 0.16 log_10_ CFU when compared with 2.984 ± 0.45 log_10_ CFU in the spleen of control animals. Both IM NCRPT groups showed no CFU and were marked at the limit of detection (10 CFU) (**Figure 4D**). To assess RPT resistance of the bacterial colonies seen in treatment groups, these colonies were spotted on 7H11 plates containing 0.075 μg/ml RPT (the concentration was selected based on the MIC value we obtained in our drug susceptibility testing of Mtb H37Rv as 0.06 μg/ml; n=3 for all treatment groups). No growth was observed, indicating the bacteria were not resistant to RPT. Treatment efficacy is also represented by the lung pathology. Lungs isolated from the control group indicate granulomatous lesions, while those isolated from the treatment groups appear clear, corroborating lung CFU data **(Figure S8)**. We also recorded the weight of the lungs and spleens isolated from animals. Lung weight data showed significantly higher weights for the untreated group, indicating inflammation caused due to bacterial infection. All rifapentine treatment regimens resulted in significantly lower lung weights **(Figure S9)**. No significant difference was obtained between the control and treatment groups for spleen weight **(Figure S9)**. Similar results were obtained for a shorter-duration independent experiment of the pre-exposure prophylactic mouse model, which consisted of a week of treatment to compare the efficacy of NCRPT formulation with daily oral dosing (**Figure S10**). We also compared the efficacy of administration of once-a-week RPT oral 20 mg/kg dose with that of NCRPT IM 60 mg/kg. It was observed that 60 mg/kg NCRPT administered intramuscularly once a week performed significantly better than 20 mg/kg RPT given orally once a week **(Figure S11)**, demonstrating the effect of sustained-release nanocrystal RPT through IM injection.

**Figure 4.**
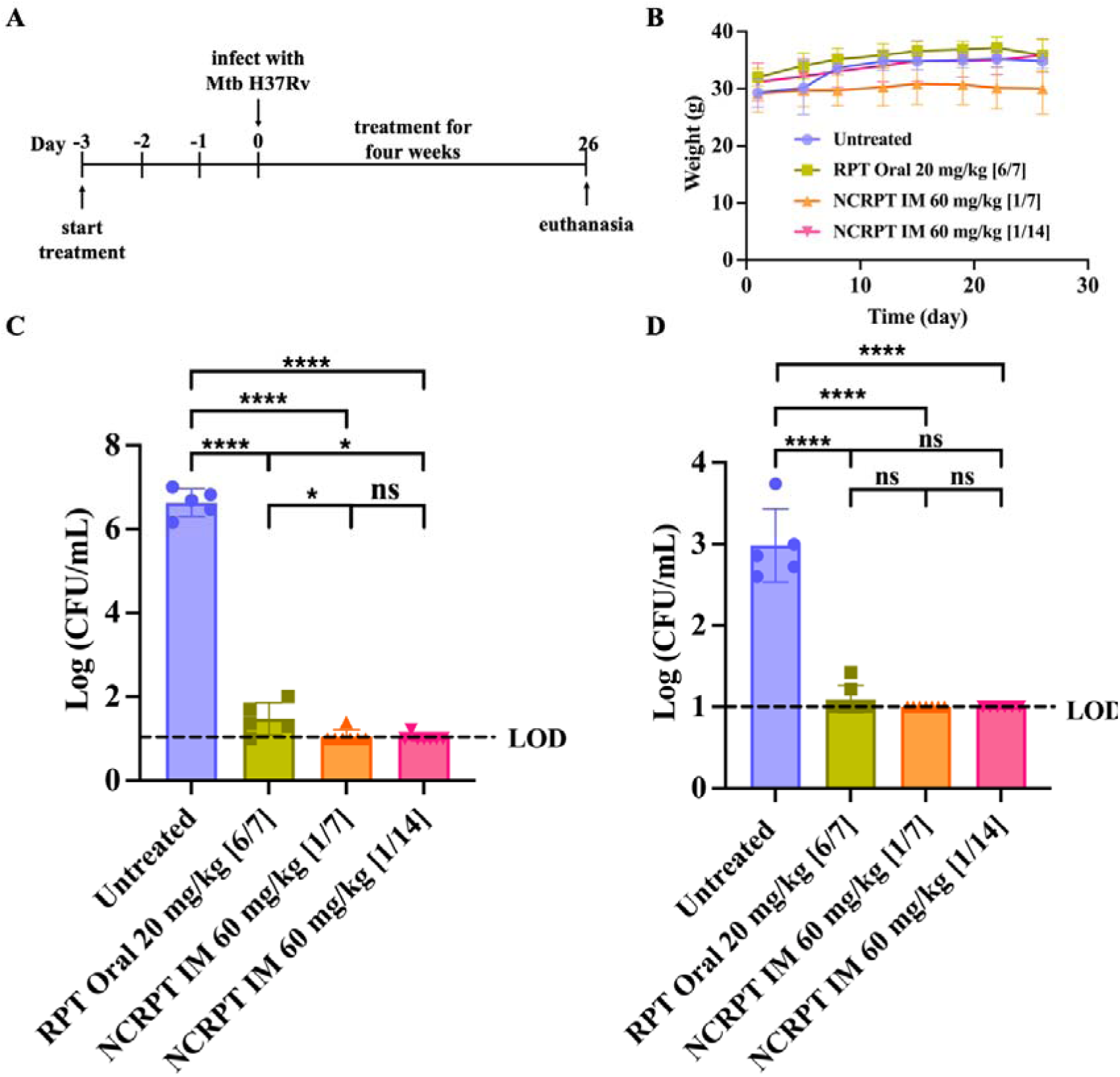
Pre-exposure prophylaxis efficacy of NCRPT in mice infected with Mtb H37Rv. (**A**) Experiment scheme and treatment regimen. The first dose of treatment was administered before the Mtb infection. Treatment was given for a month. (**B**) Weight chart of mice. Weights were checked twice a week for four weeks (n=5 distinct animals for control, n=7 distinct animals for all the other groups). Figures in square parentheses represent dosage frequency. [6/7] represents drug administration 6 days a week, [1/7] represents drug administration once a week and [1/14] represents drug administration once every two weeks. (**C**) Bacterial count in the lung on day 26 after treatment started was plotted as Log CFU/mL (n=5 lung samples from distinct animals for the control and RPT oral 20 mg/kg group, n=7 lung samples from distinct animals for both the NCRPT IM groups). (**D**) Bacterial count in spleen on day 26 after treatment started plotted as Log CFU/mL (n=5 spleen samples from distinct animals for control and RPT oral 20 mg/kg group, n=7 spleen samples from distinct animals for both the NCRPT IM groups). Data in the graph represents mean ± SD. For **C** and **D**, p-values were determined by ordinary one-way ANOVA and Tukey’s multiple comparison tests. p-value < 0.05 was considered significant. **** p< 0.0001 for untreated vs all treatment groups). *p = 0.04 for RPT Oral vs NCRPT IM [1/7], *p = 0.028 for RPT Oral vs NCRPT IM [1/14] in **C**. ns = non-significant. LOD = limit of detection.

### Efficacy of NCRPT in a therapeutic mouse model

After testing the efficacy of injectable RPT nanocrystal formulations in a pre-exposure prophylactic animal model, we further tested the efficacy of this formulation as a proof-of-concept in a therapeutic TB model. The schematic of the experimental design is detailed in **Figure 5A**. Animals were infected with Mtb H37Rv, and the infection was allowed to develop for four weeks, followed by four weeks of treatment. Treatment groups included either orally administered monotherapy of RPT 6 days a week or NCRPT formulation administered intramuscularly at a frequency of once a week and once every two weeks. Animal weights recorded at regular intervals throughout the treatment duration showed no weight loss, indicating the safety of the formulations **(Figure 5B)**. The day after infection, the mean log_10_ CFU in the lung was 3.15 ± 0.13. Four weeks post-infection, at the start of the treatment, the mean log_10_ CFU in the lung increased to 6.186 ± 0.32. Bacterial load was assessed in both the lung and spleen after two weeks of treatment and after four weeks upon completion of treatment. Two weeks post treatment, all the treatment groups showed reduced bacterial load in lungs of 4.367 ± 0.34, 4.23 ± 0.33 and 4.29 ± 0.35 for oral, NCRPT (1/7) and NCRPT (1/14) groups, respectively, when compared with the control group (infected, untreated) that harbored a bacterial load of 5.95 ± 0.26 log_10_ CFU **(Figure 5C)**. A similar reduction in bacterial load was observed upon four weeks of treatment. The mean log_10_ CFU in the lung was observed to be 1.95 ± 0.68, 2.36 ± 0.75 and 2.88 ± 0.62 for oral, NCRPT (1/7) and NCRPT (1/14) groups, respectively, when compared with the control group (infected, untreated) that harbored a bacterial load of 6.11 ± 0.53 log_10_ CFU **(Figure 5D)**. A significant reduction was observed in the spleen bacterial load of the treated group compared to the infected untreated control. Two weeks post-treatment, the mean log_10_ CFU in the spleen of control animals was observed to be 2.567 ± 0.537, while that of the treated groups was found to be 1, 1.01 ± 0.05, 1.49 ± 0.51 in the oral, NCRPT (1/7) and NCRPT (1/14) groups, respectively **(Figure 5E)**. Upon completion of four weeks of treatment, bacterial load in the spleen for treated groups was observed to be at the limit of detection (LOD) of mean log_10_ CFU 1, 1.01 ± 0.05 and 1 for oral, NCRPT (1/7) and NCRPT (1/14) groups, respectively. Bacterial load in the spleen for control animals was a mean log_10_ CFU of 2.35 ± 0.41 **(Figure 5F)**. No statistical difference was observed for both lung and spleen CFU counts between the treatment groups. To assess RPT resistance of the bacterial colonies seen in treatment groups, these colonies were spotted on 7H11 plates containing 0.075 μg/ml RPT (n=5 for all treatment groups). No growth was observed, indicating the bacteria are not resistant to RPT. Lung weight data showed significantly higher weights for the untreated group, indicating inflammation caused due to bacterial infection. All rifapentine treatment regimens resulted in significantly lower lung weights. No significant difference was obtained between the control and treatment groups for spleen weight **(Figure S12)**. Treatment efficacy is also represented by the lung pathology. Lungs isolated from the untreated group showed granulomatous lesions, while those isolated from the treatment groups appeared clearer, corroborating lung CFU data both after two **(Figure S13)** and four weeks of treatment **(Figure S14)**.

**Figure 5.**
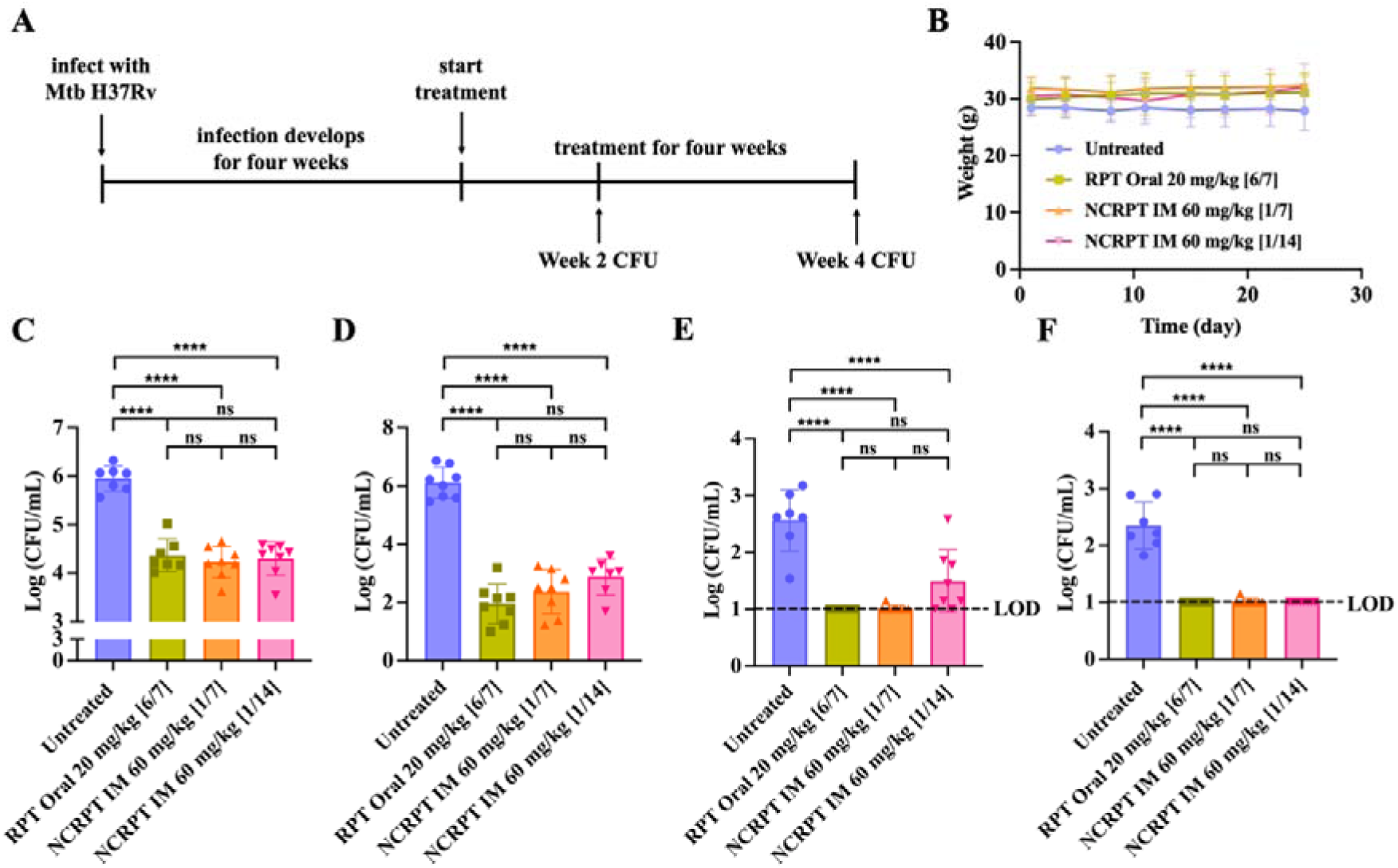
Therapeutic efficacy of NCRPT in mice infected with Mtb H37Rv. (**A**) Experimental scheme and treatment regimen. Treatment began four weeks post-infection and continued for a month. (**B**) Weight chart of mice. Weights were checked twice a week for four weeks (n=8 distinct animals for all groups). Figures in square parentheses represent dosage frequency. [6/7] represents drug administration 6 days a week, [1/7] represents drug administration once a week and [1/14] represents drug administration once every two weeks. (**C**) Bacterial count in the lung two weeks post-treatment plotted as Log CFU/mL (n=7 lung samples from distinct animals for control and n=8 lung samples from distinct animals for all the other groups). (**D**) Bacterial count in the lung four weeks post-treatment plotted as Log CFU/mL (n=8 lung samples from distinct animals for all groups). (**E**) Bacterial count in spleen two weeks post-treatment plotted as Log CFU/mL (n=7 spleen samples from distinct animals for control and n=8 spleen samples from distinct animals for all the other groups). (**F**) Bacterial count in spleen four weeks post-treatment plotted as Log CFU/mL (n=8 spleen samples from distinct animals for all groups). Data in graphs for **B**, **C, D, E** and **F** represent mean ± SD. p values were determined by ordinary one-way ANOVA and Tukey’s multiple comparison tests. p-value < 0.05 was considered significant. **** p< 0.0001 for untreated vs all treatment groups. ns = non-significant. LOD = limit of detection.

## Discussion

In this study, we developed an injectable nanocrystalline formulation of rifapentine (NCRPT), a first-line anti-TB drug, to reduce the dosage frequency of RPT administration and improve patient compliance for TB treatment. RPT has been approved for a one-month daily oral regimen along with isoniazid for TB preventive treatment. Six weeks of daily RPT alone has been shown to be equally effective as one month of daily oral RPT and isoniazid and three months of weekly oral RPT and isoniazid for latent TB infection treatment [29]. Our goal was to develop a formulation to reduce the dosage frequency while maintaining serum RPT concentration above 2 μg/mL [28]. It is well known that the amorphous nature of the drug formulation leads to higher dissolution of the drug owing to its higher internal energy and molecular motion compared to the crystalline formulation [30]. The amorphousness of the drug in a formulation is particularly of interest as it leads to higher bioavailability for poorly water-soluble drugs administered orally [31]. The intrinsic long-range order and stability of crystalline drugs can lead to slower dissolution of the drug from dissolution-controlled drug depots. This effect has been demonstrated by Appell *et al.* for long-acting moxifloxacin injectable formulation [23]. In our study, we developed long-acting crystalline and amorphous formulations of RPT (**Figure 1A-C**) and compared the drug dissolution both in an *in vitro and in vivo* setup. Characterization using diffraction (pXRD) and thermal analysis (DSC) clearly indicated the difference in the nature of these formulations (**Figure 1D-I**) in accordance with the published guidelines [30]. *In vitro* release assay clearly demonstrates the faster dissolution of RPT from ARPT compared to both the crystalline formulations (**Figure 2A**). As the *in vitro* setup is a static model and does not recapitulate the effect of interstitial flow in dissolution of the drug, we tested the drug dissolution from depots in an *in vivo* model. Mice tissue analysis post-intramuscular injection indicated that the depot formed from the crystalline formulation of rifapentine (NCRPT) had ∼7-fold higher retention of RPT in the muscle compared to the amorphous formulation on day 5 (**Figure 2B**). To the best of our knowledge, this is the first study to highlight the advantage of crystalline formulations over amorphous solid dispersions in forming *in vivo* drug depots with slower dissolution for an intramuscularly administered long-acting injectable. It indicates the potential of using crystallization as a technique in developing LAIs with reduced dosage frequency for better patient compliance. To our surprise, we do not observe any difference in drug dissolution between NCRPT and MCRPT in either the *in vitro* or the *in vivo* models suggesting that crystallinity plays a major role in controlling dissolution rates compared to size of the particles. Given the different techniques involved in preparting amorphous and crystalline formulations, the use of different surfactants was unavoidable. However, the extremely high loading levels of rifapentine suggests that surfactants are unlikely to be a major contributor to the differences observed in dissolution rates. In addition, surfactants are only expected to be on the surface of the crystals which are likely to be solubilized as surface of particles dissolves. Additionally, the crystals in the MCRPT formulation tend to settle faster compared to NCRPT, rendering the formulation non-ideal for injections. NCRPT formulations exhibited high stability and injectability (Figure S3 and SEM size analysis). Since the nanocrystalline formulation of RPT was found to be more suitable for our purpose of developing RPT LAI, a pharmacokinetic study was performed with NCRPT. We found that NCRPT resulted in sustained release of therapeutically required concentrations for 6-9 days with one administration of IM injection in mice model (**Figure 3B**). This pharmacokinetic profile could be the combined result of the hydrophobicity of RPT, the crystalline nature of the injected formulation, and the IM route of administration. Together, these factors lead to the formation of a dissolution-controlled drug depot at the site of administration, which gradually dissolves and steadily releases the required amounts of drug for several days. To verify whether these sustained levels of RPT would lead to antimicrobial efficacy, we tested our formulation on a pre-exposure prophylactic TB-infected mice model. Our results indicated that when NCRPT was administered intramuscularly once a week at a dose of 60 mg/kg, it performed equally well as that of orally administered RPT given six days a week at a dose of 20 mg/kg (**Figure 4**). This demonstrated the potential of injectable crystalline formulations in achieving extended release of therapeutically required drug levels, consequently reducing dosage frequency. It is possible that the reduction in bacterial load observed for once in two weeks administration of NCRPT in our pre-exposure prophylactic model could be the result of exposing the animals to the drug prior to bacterial infection, which is enough to eliminate most of the bacteria at the time of infection (3 days post CRPT injection). Hence, as a proof of concept, we next tested our formulations in a therapeutic regimen in the TB mice model. In these experiments, the animals were infected with Mtb for a month before starting drug treatments. We observed that 60 mg/kg NCRPT formulation administered intramuscularly at either once a week or once every two weeks performed equally well as orally administered RPT given six days a week at a dose of 20 mg/kg (**Figure 5**).

Chang *et al.* conducted a study with oral dosing of RPT to simulate exposure profiles of long-acting formulations for TPT [28]. RPT was orally administered twice daily at low doses to achieve trough plasma concentrations of 0.18, 0.6, 2 and 3.5 μg/mL. It was observed that maintaining plasma concentration of 0.18 μg/mL or 0.6 μg/mL did not have any bactericidal effect. They also mimicked LAI formulation pharmacokinetics by giving multiple injections every day and achieved RPT concentrations falling from a peak of 2 μg/mL to the trough concentration of 0.6 μg/mL in 4 weeks. This regimen showed bactericidal activity for this period. This suggests that the minimum RPT serum concentration to exhibit bactericidal activity is between 0.6 to 2 μg/mL. In our study, we observed plasma concentrations of 2.48 μg/mL on day 6 and 0.42 μg/mL on day 9 post one injection of a 60 mg/kg dose, which could account for the efficacy we observed with once-a-week administration of NCRPT. To our surprise, we observed that NCRPT, when administered once every two weeks at a dose of 60 mg/kg, performed equally well as that for once a week 60 mg/kg IM administration. The bactericidal effect seen for once in two-week administration, despite low levels of RPT expected from 7-14 days after administration, is in contrast with the study mentioned. Another reason for the bactericidal activity of RPT at lower plasma levels could be attributed to the post-antibiotic effect (PAE) of RPT. PAE refers to the continuous suppression of bacterial growth followed by limited exposure to an antibiotic [32]. In a study carried out to evaluate the PAE of RPT, Mtb H37Rv was exposed to 1 μg/mL of RPT for different durations [33]. It was observed that a 4-day exposure of RPT at 1 μg/mL led to the slower growth rate of the bacteria for 5 days, beyond which the growth rate increased. A similar PAE effect was observed in humans [34]. We speculate that in our case, the bactericidal activity of once in two weeks NCRPT administration could be due to the PAE effect of RPT beyond day 6 until the next dose is administered at day 14. We also did not observe any emergence of RPT resistance in any of our animal studies, though more extensive analysis is required in future studies. Efficacy of our formulation against clinical strains of Mtb remains to be tested, which could have different susceptibility towards RPT.

The potential of LAI formulation in reducing dosage frequency for TPT has been demonstrated previously. A long-acting injectable formulation of a novel compound targeting purine synthesis in Mtb exhibited four weeks of sustained release upon subcutaneous injection in mice model considerably reducing dosage frequency [35]. In a recent study, self-assembling peptide hydrogel has been demonstrated to provide controlled release of ganfeborole (a molecule in TB clinical trial) over 14 days [36]. Intramuscularly administered bedaquiline LAI formulations have also been developed [19]. This study tested several regimens, including either one or two injections of bedaquiline or combining oral and injectable bedaquiline for TPT in a paucibacillary mouse model. Efficacy was observed for regimens containing either two weeks of oral bedaquiline along with one injection of 160 mg/kg bedaquiline or four weeks of oral bedaquiline along with two injections of 160 mg/kg bedaquiline. We did not test the efficacy of combining oral and NCRPT IM administration, which can be tested in future studies and could further improve the outcomes. Studies carried out to assess preferences for TB preventive therapy have shown an increasing number of patients preferring weekly oral administration of RPT with isoniazid over daily administration. Reasons for this preference include a lesser pill burden, fewer medication days and reduced side effects due to reduced frequency of drug administration [37,38]. Further strategies and innovations are required to extend the release of RPT to over a month to further reduce the dosage frequency and make it patient-compliant. Developing similar injectable formulations for other first-line anti-TB drugs and the development of a single IM injection comprising all first-line anti-TB drugs has the potential to revolutionize TB treatment.

In summary, crystalline formulations show a much longer release of drugs from dissolution-controlled depots compared to amorphous formulations. IM administration of nanocrystalline rifapentine once a week or once every two weeks showed similar efficacy as daily oral administration of RPT. Such regimens have the potential to reduce pill burden and have fewer medication days. This proof-of-concept study could provide key insights into the development of future LAIs. It also opens an avenue into reducing dosage frequency and improving patient compliance for TB treatment by developing injectable crystalline drug formulations.

## Acknowledgements

The authors recognize the Advanced Facility for Microscopy and Microanalysis (AFMM) at IISc for access to scanning electron microscopy and the Solid State and Structural Chemistry Unit (SSCU) Central Facility at IISc for access to the powder XRD facility. The authors recognize S. Bose, Professor, Department of Materials Engineering, IISc, for access to the DSC instrument and help with analysis. We also acknowledge the Biosafety Level 3 (BSL-3) facility at the Centre for Infectious Diseases and the Central Animal Facility at IISc. The following funding sources are also acknowledged:

Scheme for Translational and Advanced Research-Ministry of Education (MoE-STARS/STARS-2/2023-0329) research funding to Rachit Agarwal.

Indian Institute of Science IoE postdoctoral fellowship (R(HR)(loE-IISc)(PDF)(BSSE)(RS)-591) to Rituparna Saha.

Prime Minister’s Research Fellowship (PMRF) to Nisha Sanjay Barge granted by the Ministry of Education, Government of India (PMRF ID - 0202578).

Prime Minister’s Research Fellowship (PMRF) to Yeswanth Chakravarthy Kalapala, granted by the Ministry of Education, Government of India (PMRF ID - 0201550).

## Author Contribution Statement

Conceptualization: RA, NSB

Methodology: RA, NSB

Investigation: NSB, YCK, PR, AAD, NKB, SV

Supervision: RA

Writing – original draft: RA, NSB

Writing – review and editing: RA, NSB, RS, HC

## Data availability

All data supporting the findings of this study are available within the article and its Supplementary Information file.

## Materials and methods

### Study approvals

All experiments with Mtb H37Rv were approved (IBSC/IISc/RA/03/2022-2023) by the Institutional Biosafety Committee (IBSC), Indian Institute of Science (IISc) and carried out in BSL-3 laboratory settings at the Centre for Infectious Diseases, IISc. All animal experiments were approved (CAF/Ethics/951/2023) by the Institutional Animal Ethics Committee (IAEC), IISc.

### Synthesis and characterization of RPT formulations and dissolution of RPT formulations

#### Synthesis of crystal RPT formulations

Rifapentine (RPT) was purchased from TCI Chemicals, Tokyo. For crystal RPT formulations, RPT was crystallized using the solvent-antisolvent precipitation technique. RPT at a concentration of 10 mg/mL was dissolved in acetone until a clear solution was obtained. Hexane is added to the solution in a solvent: anti-solvent ratio of 1:5. The reaction is then incubated at 15 °C for 20 hours. The reaction leads to the formation of RPT crystals. The crystals are dried overnight in an incubator at 37 °C. For NCRPT, a 5 mg/mL suspension of RPT crystals (in Milli-Q ultrapure water) containing 1.5 mg/mL Poloxamer P407 (Sigma-Aldrich, 16758) surfactant (to avoid aggregation of nanocrystals) was used for ultrasonication. These crystals were ultrasonicated to obtain nanocrystals. Ultrasonication was carried out using a probe sonicator (QSonica sonicator, Q125) at 85 % amplitude for 5 min. Post-sonication, the suspension was washed with Milli-Q ultrapure water to remove excess surfactant, then lyophilized to remove any residual solvent and stored at room temperature for further use.

For MCRPT formulation, RPT crystals were bath sonicated in Milli-Q ultrapure water at a concentration of 10 mg/mL for 3 mins. These crystals were then allowed to settle for 5 minutes, and the supernatant was removed and replaced with fresh water. The process was performed twice. The microcrystals were lyophilized and stored at RT for further use.

#### Synthesis of ARPT formulation

RPT was dissolved in dichloromethane (DCM) at a concentration of 25 mg/mL until a clear solution was obtained. 1 % polyvinyl alcohol (PVA) (Sigma-Aldrich, average Mw 31,000 – 50,000, 363073) in water was added in the ratio of 1:4 RPT-DCM: PVA in a total of 10 mL solution. The solution is homogenized at 3000 rpm for 2 min using IKA T18 digital ULTRA TURRAX^®^ homogenizer. DCM from the suspension was allowed to evaporate, post which the suspension was washed with Milli-Q ultrapure water and lyophilized. The lyophilized formulation is used for further characterization.

#### Scanning Electron Microscopy and size quantification

RPT crystals, RPT nanocrystals, RPT microcrystals and ARPT were imaged using ThermoFisher Apreo 2S HiVac Schottky Field Emission Scanning Electron Microscope (FESEM). MCRPT formulation was directly placed on carbon tape on a specimen stub before gold coating. RPT crystals, NCRPT and ARPT were suspended in Milli-Q ultrapure water at a concentration of 1 mg/mL for crystals. 10 μL of these suspensions were put onto a carbon tape on a specimen stub. The samples were dried for 48 hours under vacuum before imaging. The sample was gold sputter-coated for 30 seconds and imaged under high vacuum at 5 kV. The size of the nanocrystals in NCRPT (n = 300) from three distinct batches, microcrystals in MCRPT (n = 306) from three distinct batches and particles in ARPT (n = 250) from one batch were quantified using ImageJ software.

#### Powder XRD and DSC measurements

Powder XRD measurements for NCRPT, MCRPT and ARPT were carried out using the Bruker D8 Advance X-ray diffractometer. The diffracted pattern was recorded for 2θ values from 5 to 60°. A copper (Cu) X-ray source was used, which produces *Kα* radiation at a wavelength of ∼ 1.54 Å. A Nickel (Ni) filter is used to selectively absorb the *Kβ* wavelength. The instrument operates in Bragg-Brentano geometry where the X-ray source, sample, and detector lie on a focusing circle. The sample rotates at an angle of θ, and the detector moves at 2θ. Differential scanning calorimetry measurements for NCRPT, MCRPT and ARPT were carried out using TA Instruments DSC Q2000. Standard DSC measurements were performed at a heating rate of 10 °C/min.

#### Loading levels of RPT formulations

To quantify RPT loading levels in NCRPT and MCRPT, the formulations processed from three different batches were analyzed. NCRPT and MCRPT 1 mg/mL stock solution was prepared in methanol. The solution was diluted to 100 μg/mL in methanol. Absorbance of the solution was read at 486 nm (maximum absorbance of RPT determined in a separate assay; results not included). The concentration of the solution was quantified using a standard curve of RPT in methanol. Loading level of RPT was quantified using the following equation: (Concentration from absorbance/100 μg/mL) *100 %

To quantify RPT loading levels in ARPT, ARPT formulations processed from three different batches were analyzed. ARPT 1 mg/mL stock solution was prepared in DMSO. The solution was diluted to 100 μg/mL in DMSO. Absorbance of the solution was read at 486 nm (maximum absorbance of RPT determined in a separate assay; results not included). The concentration of the solution was quantified using a standard curve of RPT in DMSO. Loading level of RPT was quantified using the following equation: (Concentration from absorbance/100 μg/mL) *100 %

### RPT crystal dissolution

RPT crystals were incubated at a concentration of 0.01 mg/mL in 1X Phosphate-buffered saline (PBS) in a 96-well plate. One crystal was selected from each well (total 9 wells) and repeatedly imaged by fixing the selection pane in IN Cell Analyzer 6000 (GE Healthcare) brightfield microscope at regular intervals for a total of 24 hours. The crystal area was calculated from the images using ImageJ software by tracing the crystal boundaries.

### RPT MIC determination with Mtb H37Rv

Resazurin microtiter plate assay was used to determine the MIC of RPT. Drug concentrations ranging from 0.15 μg/mL to 960 μg/mL with two-fold increasing intervals were tested. 1 mg/mL of RPT stock was prepared in DMSO. Subsequent dilutions of the drug were prepared in 7H9 media (supplemented with ADC). 100 μL of the drug solution was incubated with 100 μL of bacterial culture (density of 1 × 10^6^ CFU/mL) in a 96-well plate for 7 days. Post drug incubation, 30 μL of 0.01 % resazurin salt (resazurin sodium salt, Sigma, R7017) solution was added, and the plate was incubated at 37 °C for 24 hours. Post incubation, the plate was read through fluorescence with excitation at 530 nm and emission at 590 nm. The lowest concentration to inhibit bacterial growth was determined as the MIC.

### *In vitro* dissolution study with NCRPT, MCRPT and ARPT

The *in vitro* dissolution study was performed using dialysis units (Thermo Scientific™ Slide-A-Lyzer™ Mini Dialysis Devices, 10K MWCO). 1 mg/mL of NCRPT, MCRPT and ARPT formulations in 1X PBS were added to the 0.5 mL dialysis cups. The cups were suspended in 15 mL conical tubes filled with 14 mL 1X PBS (PBS was changed every 24 h to simulate an infinite sink condition). Samples from dialysis cups were analyzed by dissolving in methanol to quantify RPT amount every 24 h by recording absorbance at 486 nm.

### *In vivo* depot study with NCRPT, MCRPT and ARPT

NCRPT, MCRPT and ARPT suspensions in 1X PBS were administered to male C57BL/6 mice at a dose of 60 mg/kg via a single IM injection in the caudal thigh muscle using a 27 G × ½” needle. Mice were euthanized within 2 hours post-dosing (Day 0) and 120 hours post-dosing (Day 5). The caudal thigh muscle at the site of injection was isolated in 1 mL methanol for the NCRPT and MCRPT groups and 1 mL DMSO for the ARPT. RPT from the depot site was allowed to dissolve in either methanol or DMSO by incubating for 24 hours at 4 °C. Post incubation, the absorbance of the sample was recorded at 486 nm, and the RPT amount at the depot site was quantified using standard curves made in either methanol or DMSO.

### Injectability and stability of NCRPT formulation

NCRPT formulation was prepared at a concentration of 15 mg/mL (corresponding to a 60 mg/kg dose). The RPT concentration in this formulation was determined by dissolving the NCRPT formulation in methanol and reading the absorbance at 486 nm to quantify RPT using a standard curve made in methanol. The formulation was passed through a 27 G × ½” needle. The RPT concentration in this formulation was determined by dissolving the NCRPT formulation in methanol and reading the absorbance at 486 nm to quantify RPT using a standard curve made in methanol. This assay was performed on three distinct batches of NCRPT.

The size of the nanocrystals was determined before and after injection using SEM. 10 μL of NCRPT formulation at a concentration of 15 mg/mL (corresponding to a 60 mg/kg dose) was loaded onto a carbon tape on a specimen stub before gold coating. The formulation was passed through a 27 G × ½” needle. 10 μL of this formulation was loaded onto a carbon tape on a specimen stub before gold coating. The samples were dried for 48 hours under vacuum before imaging. The sample was gold sputter-coated for 30 seconds and imaged under high vacuum at 5 kV. The size of the nanocrystals in NCRPT before injection (n = 300) from three distinct batches and post-injection (n = 306) from three distinct batches was quantified using ImageJ software.

### *In vitro* cytotoxicity of CRPT formulations

Cytotoxicity of NCRPT formulations was assessed with THP-1 macrophages (obtained from National Centre for Cell Science, Pune) by WST-1 cell proliferation assay using a kit-based method (Cayman Chemicals, 10008883). THP-1 monocytes were seeded into a 96-well plate at a density of 2×10^4^ cells/well and differentiated into THP-1 macrophages using phorbol 12-myristate 13-acetate (PMA) (Sigma, P1585) treatment (20 ng/mL). A stock of 20 mg/mL commercially available RPT was made in DMSO. For CRPT, RPT nanocrystals were dissolved in 1X PBS to make a stock of 1 mg/mL. Cytotoxicity for both formulations was assessed at varying concentrations of 100 μg/mL, 30 μg/mL, 3 μg/mL, 0.3 μg/mL, 30 ng/mL and 3 ng/mL. Differentiated THP-1 macrophages were incubated with the above mentioned concentrations of either RPT or CRPT for 24 hours. Post incubation, WST reagent was added to the wells, allowed to incubate for 2 hours, and the plate was read at 450 nm.

Cytotoxicity of NCRPT formulations was also assessed with the human embryonic kidney (HEK-293) cell line. Cells were seeded into a 96-well plate at a density of 1.2×10^4^ cells/well and incubated at 37 °C for 24 hours. Cytotoxicity for both formulations (NCRPT and commercially available RPT) was assessed at varying concentrations of 30 μg/mL, 3 μg/mL, 0.3 μg/mL, 30 ng/mL. Drug formulations were incubated with the cells for 24 hours. Post incubation, WST reagent was added to the wells, allowed to incubate for 4 hours, and the plate was read at 450 nm.

### *In vitro* efficacy of NCRPT formulations

*In vitro* efficacy of NCRPT formulation was assessed with THP-1 macrophages infected with Mtb H37Rv to test whether the crystallization process affects the antibacterial efficacy of RPT. THP-1 monocytes were seeded into a 96-well plate at a density of 2×10^4^ cells/well and differentiated into THP-1 macrophages using PMA treatment (20 ng/mL). Subsequent protocol was carried out in BSL-3 settings. THP-1 macrophages were then infected with Mtb H37Rv mid-log secondary culture with OD ∼ 0.6-0.8. The infection was allowed to develop for 4 hours. Post infection, the infected cells were treated with 0.2 mg/mL filter-sterilized amikacin (HiMedia, CMS644) solution prepared in RPMI media for 2 hours to eliminate extracellular bacteria. Post amikacin treatment, infected macrophages were either treated with 5 μg/mL free RPT (solution prepared by dissolving RPT in 1X PBS containing 5 % DMSO) or 5 μg/mL NCRPT (solution prepared by dissolving NCRPT in DMSO at a 1 mg/mL concentration. Subsequent dilution was made in 1X PBS) for 48 hours. Untreated cells were considered the control. Post treatment, cells were lysed by treating with 0.1 % filter-sterilized Triton-X (Thermo Scientific Chemicals) solution for 10 min. Cell lysate was plated on Middlebrook 7H11 agar base (HiMedia, M511) plates containing Middlebrook OADC growth supplement (HiMedia, FD018) and incubated at 37 °C to quantify bacterial load.

### Serum pharmacokinetics of NCRPT

**The** pharmacokinetic study of NCRPT was carried out in male Balb/c mice (> 20 g, 6-8 weeks). Animals were administered 60 mg/kg of CRPT formulation (100 μL of 15 mg/mL solution) via a single IM injection in the caudal thigh muscle using a 27 G × ½” needle. Blood from these animals was collected via retro-orbital bleeding at regular intervals using micro hematocrit capillary tubes. Blood samples were kept at 4 °C for 8 hours, followed by centrifugation at 8000×g for 4 min at 4 °C to collect serum. Serum was analyzed using LC-MS to quantify RPT levels. Pharmacokinetic parameters for NCRPT formulation were determined using non-compartmental analysis from serum drug concentration and time profile. Peak serum drug concentrations (C_max_) and time to achieve C_max_ (T_max_) were obtained by visual inspection of the average concentration-versus-time profiles. The area under the serum concentration-time curve from time zero to 9 days was determined by using the linear trapezoidal rule. The terminal elimination rate constant (λ_z_) was estimated by the linear least-squares regression of the log serum drug concentration-time data from the terminal elimination phase. AUC from time zero to infinity (AUC_0–∞_) was estimated by dividing the last drug concentration by λ_z_. The T_1/2_ was calculated from the equation, T_1/2_ = 0.693/λ_z_. Clearance was calculated from the equation, clearance = total dose/AUC_0-∞_. Volume of distribution was calculated from the equation, volume of distribution = clearance/ λ_z_.

### Bronchoalveolar lavage fluid (BALF) pharmacokinetics of NCRPT

Male Balb/c mice (> 20 g, 6-8 weeks) were administered with RPT orally at 20 mg/kg via oral gavage or NCRPT formulation at 60 mg/kg via a single IM injection in the caudal thigh muscle using a 27 G × ½” needle. Mice were humanely euthanized 24 hours post-dosing, and BALF was collected by flushing the lungs with 1 mL 1X PBS. BALF was analyzed using LC-MS to quantify RPT levels.

### NCRPT formulation toxicity analysis

For local toxicity analysis of IM administration of NCRPT formulation, male Balb/c mice (> 20 g, 6-8 weeks) were administered with 60 mg/kg of NCRPT formulation via a single IM injection in the caudal thigh muscle using a 27 G × ½” needle. Post euthanasia, the muscle at the site of injection was retrieved 20 mins (Day 0), one week (Day 7) and four weeks (Day 27) post-injection. Muscle tissues were fixed in 4 % paraformaldehyde upon retrieval, incubated for 2 hours at room temperature and transferred to 15 % sucrose solution for overnight incubation at 4 °C. Post-incubation, the tissues were transferred to 30 % sucrose until used for cryo-sectioning. Processed muscle tissues were sectioned into 40 μm sections (Leica cryotome CM1520) and stained using hematoxylin and eosin staining.

#### Systemic toxicity of NCRPT

To assess systemic toxicity of NCRPT, male Balb/c mice (> 20 g, 6-8 weeks) were administered with RPT orally at 20 mg/kg via oral gavage or NCRPT formulation at 60 mg/kg via a single IM injection in the caudal thigh muscle using a 27 G × ½” needle. Blood from these animals was collected via retro-orbital bleeding at day 1 and 6 post drug administration using micro hematocrit capillary tubes. Blood samples were kept at 4 °C for 8 hours, followed by centrifugation at 8000×g for 4 min at 4 °C to collect serum. Serum from untreated animals was collected similarly. Systemic toxicity was assessed by testing the serum samples for ALT and AST using International Federation of Clinical Chemistry and Laboratory Medicine method [39], and for total bilirubin using Diazotized Sulfanilic Acid (DSA) method [40]. All the tests were performed by Rohana Veterinary Diagnostic Lab, Bengaluru, Karnataka 560094, India.

### *In vivo* efficacy of NCRPT in pre-exposure prophylactic mouse model

Efficacy of NCRPT formulations was initially tested in a pre-exposure prophylactic mouse model. In the first experiment, male Balb/c mice (> 20 g, 6-8 weeks) were dosed with either 20 mg/kg RPT every day for a week orally (to represent a clinically approved dose for TB treatment) or 60 mg/kg NCRPT intramuscularly once a week. Three days after drug administration, animals were aerosol infected with log phase secondary culture (OD ∼ 0.6-0.8) of Mtb H37Rv using the Glas-Col inhalation exposure system. Treatment efficacy was assessed based on lung CFU counts post CO_2_ asphyxiation. Lungs from these animals were isolated and homogenized in 2 mL 1X PBS per lung (Omni Tissue Homogenizer, TH-20, Omni International, USA). Lung homogenate was plated on 7H11 plates containing OADC and Penta Mix (HiMedia, FD260). The colonies were counted after 4-5 weeks of growth on the plates. In the second experiment, male Balb/c mice (> 20 g, 6-8 weeks) were dosed with either 20 mg/kg RPT orally or 60 mg/kg NCRPT intramuscularly. All doses were administered once a week. Three days after drug administration, animals were aerosol infected with log phase secondary culture (OD ∼ 0.6-0.8) of Mtb H37Rv using the Glas-Col inhalation exposure system. Treatment efficacy was assessed based on lung CFU counts. To assess the efficacy of NCRPT for one-month treatment, male Balb/c mice were administered either 20 mg/kg commercially available RPT via oral gavage six days a week or 60 mg/kg of NCRPT formulation via a single IM injection in the caudal thigh muscle using a 27 G × ½” needle either once a week or once every two weeks. Untreated animals were considered the control. Three days after drug administration, animals were aerosol infected with log phase secondary culture (OD ∼ 0.6-0.8) of Mtb H37Rv using the Glas-Col inhalation exposure system. Treatment efficacy was assessed based on lung CFU counts. Five animals were euthanized to enumerate the bacterial load 24 hours post-infection in the lungs. Lungs from these animals were isolated and homogenized in 1X PBS (Omni Tissue Homogenizer, TH-20, Omni International, USA). Lung homogenate was plated on 7H11 plates containing OADC and Penta Mix (HiMedia, FD260). The colonies were counted after 4-5 weeks of growth on the plates. For the rest of the animals, drug treatment was followed for four weeks. Post-treatment, all animals were euthanized. Lung and spleen from all animals were isolated and homogenized in 1X PBS. Lung and spleen homogenates were plated on 7H11 plates containing OADC and Penta Mix. The plates were incubated at 37 °C, and the colonies were counted after 4-5 weeks of growth on the plates. Animal weights were recorded throughout the experiment duration.

### *In vivo* efficacy of NCRPT in therapeutic mouse model

Female Balb/c mice were aerosol infected with log phase secondary culture (OD ∼ 0.6-0.8) of Mtb H37Rv using the Glas-Col inhalation exposure system. Six animals were euthanized to enumerate the bacterial load 24 hours post-infection in the lungs by the procedure mentioned above. The infection was allowed to develop for four weeks. Five animals were euthanized to enumerate the bacterial load in the lungs at the start of the treatment. Animals were treated with either administration of 20 mg/kg commercially available RPT via oral gavage six days a week (to represent a clinically approved dose for TB treatment) or 60 mg/kg of NCRPT formulation via a single IM injection in the caudal thigh muscle using a 27 G × ½” needle either once a week or once every two weeks. Untreated animals were considered the control. Treatment efficacy was assessed by quantifying lung and spleen CFU counts after two and four weeks of treatment. Two and four weeks post-treatment, animals were euthanized. Lung and spleen from all animals were isolated and homogenized in in 2 mL 1X PBS per organ. Lung and spleen homogenates were plated on 7H11 plates containing OADC and Penta Mix. The plates were incubated at 37 °C, and the colonies were counted after 4-5 weeks of growth on the plates. Animal weights were recorded throughout the experiment duration.

### Assessing RPT resistance

The Agar dilution method was used to check whether the bacterial colonies obtained from animals undergoing prophylactic and therapeutic treatments were resistant to RPT. Colonies obtained on 7H11 agar plates from the treatment groups were resuspended in 1X PBS and diluted to obtain a cell density of 10^6^ CFU/mL. 10 μL (ensuring 10^4^ CFU per spot) [41] of this solution was spotted on 0.075 μg/mL RPT 7H11 agar plates supplemented with OADC and Penta Mix. The plates were incubated at 37 °C and monitored for 8 weeks.

### Statistics

All experiments were performed on independent biological replicates. Statistical significance was determined for the control and experimental groups using one-way ANOVA with Tukey’s multiple comparisons test (alpha = 0.05), or by two-way ANOVA followed by post-hoc Tukey’s multiple comparison tests, as specified. All statistical tests used were two-sided. Outlier analysis was performed using Grubbs’ test (p < 0.05 was considered significant for outlier analysis). GraphPad (Prism) was used for all statistical analyses.\

### Ethical Clearance

All experiments were performed following the Institutional BioSafety Committee (Protocol Number: IBSC/IISc/RA/07/2025) approval. For animal studies, Institutional Animal Ethics Committee (IAEC) (Protocol Number: CAF/Ethics/951/2023) approval was obtained.

