## Supplementary Information for "Dissolution-Controlled Nanocrystalline Rifapentine Formulation for Tuberculosis Treatment"

^b^ Sanjay V.

Centre for Infectious Disease Research

Indian Institute of Science

Bengaluru, India

^c^ Harinath Chakrapani

Department of Chemistry

Indian Institute of Science Education and Research

Pune, India

This file includes:

Table S1 to S2

Figures S1 to S14

| Solvent (amount in µL) | RPT amount (mg) | Anti-solvent (amount in mL) | Temp conditions (ºC) | Crystal formation | Crystal dimensions |
| --- | --- | --- | --- | --- | --- |
| Acetone (600) | 4 | Toluene (10) | -20 | No | - |
| Acetone (600) | 4 | Xylene (10) | -20 | No | - |
| Acetone (200) | 1.33 | Hexane (10) | -20 | No | - |
| Acetone (600) | 4 | Hexane (10) | -20 | Yes | 550 -600 µm length |
| Acetone (600) | 4 | Hexane (5) | -20 | Yes | 350 - 400 µm length |
| Acetone (600) | 4 | Pentane (6) | RT | Yes | 300 - 350 µm length |
| **Acetone (1800)** | **18** | **Hexane (9)** | **15** | **Yes** | **200-400 µm length** |

***Table S1: Experimental details for crystallizing RPT.*** *The condition listed in the last row was used to obtain microcrystal RPT for further experiments.*

| Drug parameter | Value |
| --- | --- |
| C_max_ (μg/mL) | 11.15 |
| T_max_ (day) | 1 |
| AUC_0-9_ (μg.day/mL) | 33.62 |
| AUC_0-∞_ (μg.day/mL) | 35.17 |
| Terminal rate constant λ_z_ (day^-1^) | 0.33 |
| T_1/2_ (days) | 2.09 |
| Clearance (L/day) | 1.7 |
| Volume of distribution (L) | 5.14 |

***Table S2:*** ***Pharmacokinetic parameters for NCRPT.***

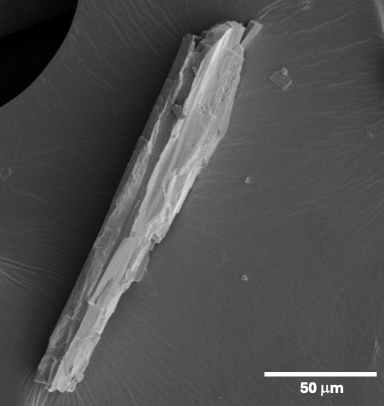

***Figure S1.*** *SEM image of RPT crystal obtained after solvent-antisolvent precipitation technique (scale bar - 50 µm).*

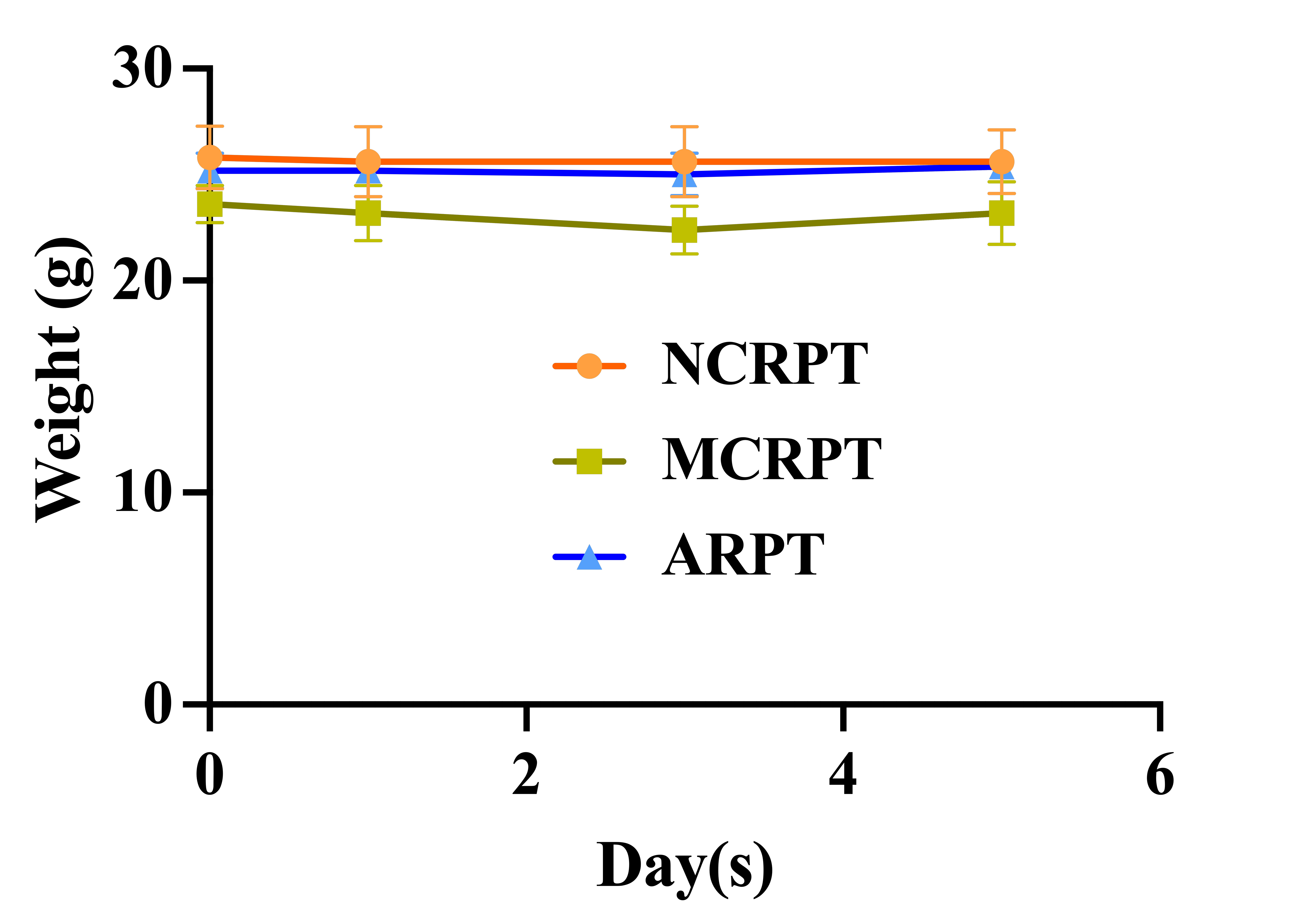

***Figure S2.*** ***Weight chart of mice for the in vivo depot study.*** *Weights were checked every other day until day 5 (n = 5 animals for both groups).*

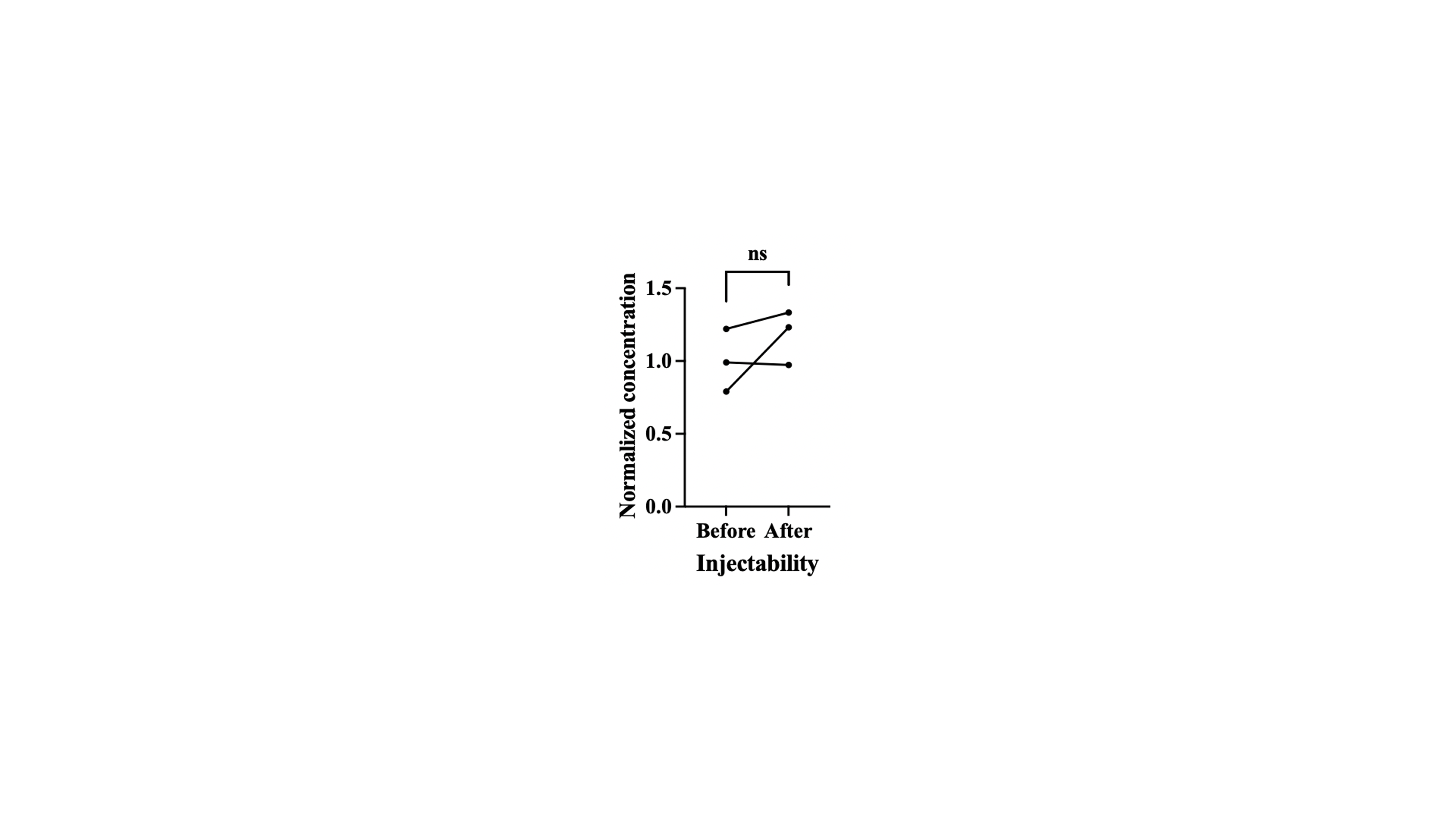

***Figure S3.*** *Injectability of NCRPT. Normalized concentration of RPT in NCRPT formulation before and after passing through 27 G × ½” needle*. *Data in the graph represent mean ± SD and was analyzed by paired t-test. p-value < 0.05 was considered signiﬁcant. ns = non-significant.*

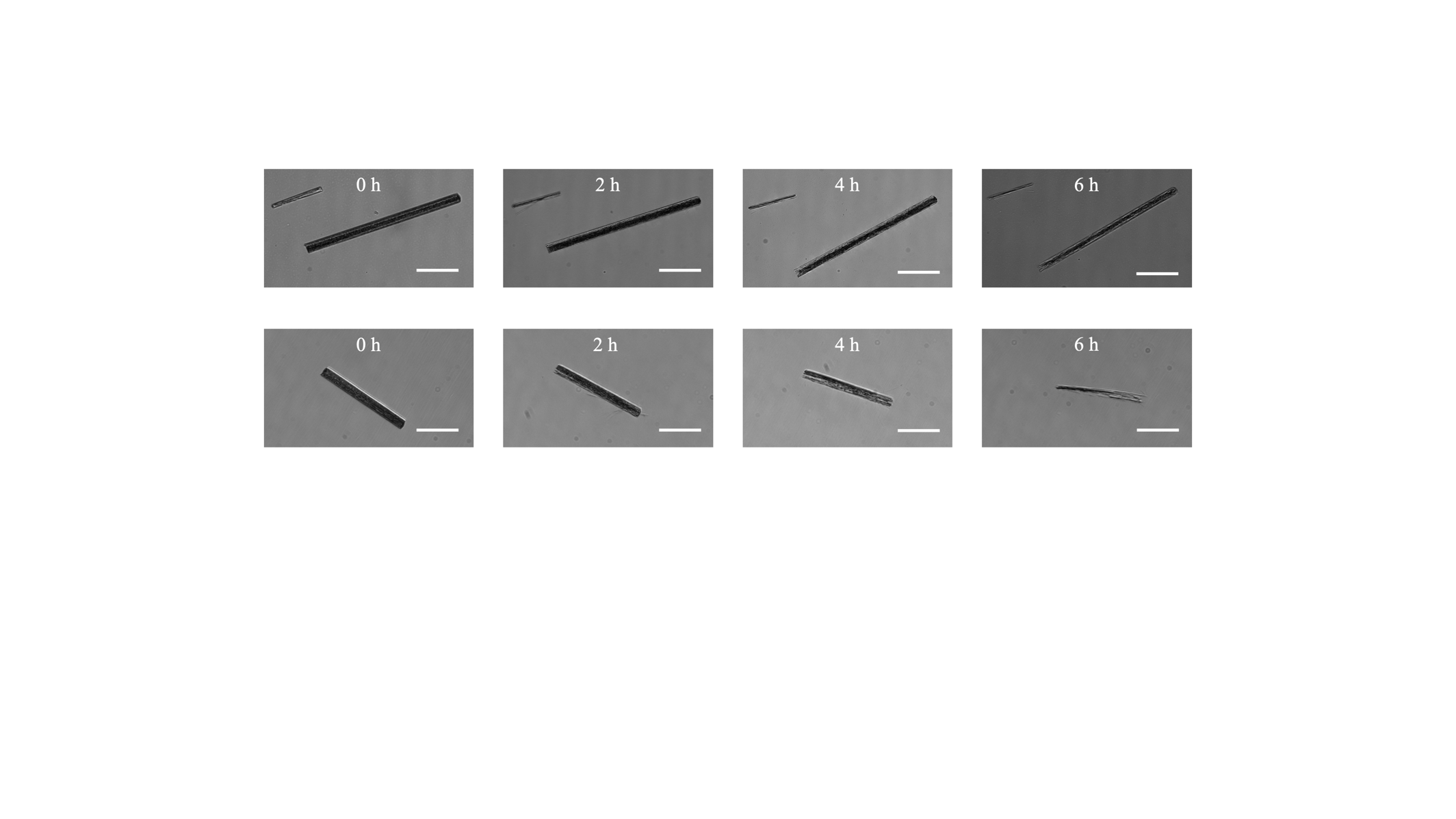

***Figure S4. Surface dissolution of crystals.*** *Representative RPT crystals at 0.01 mg/mL concentration in 1X PBS imaged over a period of 6 hours. Scale bar = 100 μm.*

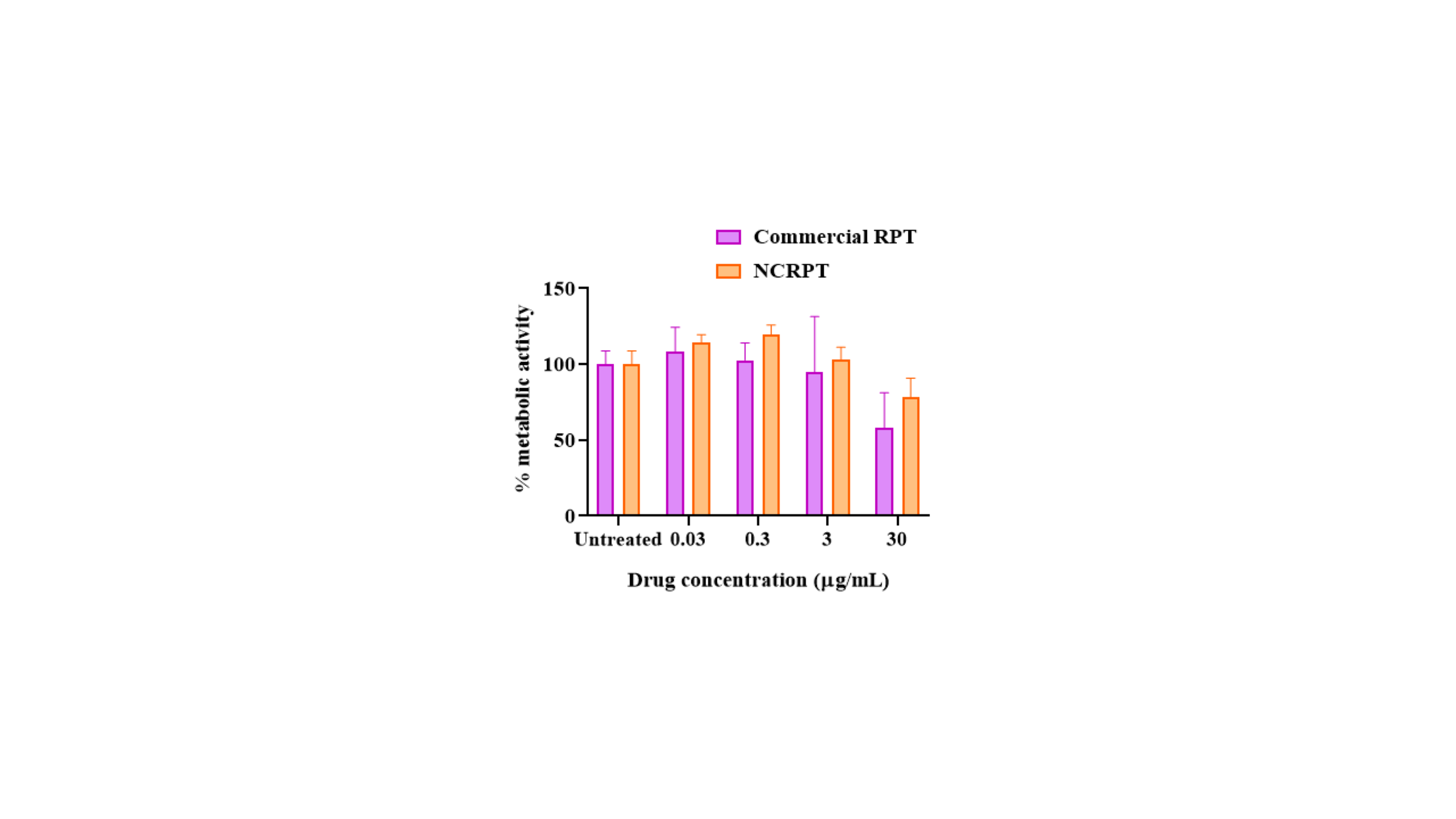

***Figure S5. NCRPT is cytocompatible with HEK-293 cells.*** *WST assay performed with HEK-293 cells to determine the cytocompatibility of NCRPT formulation compared with the commercially available rifapentine as control (n=3 distinct samples for both groups). Data in the graph represent mean ± SD.*

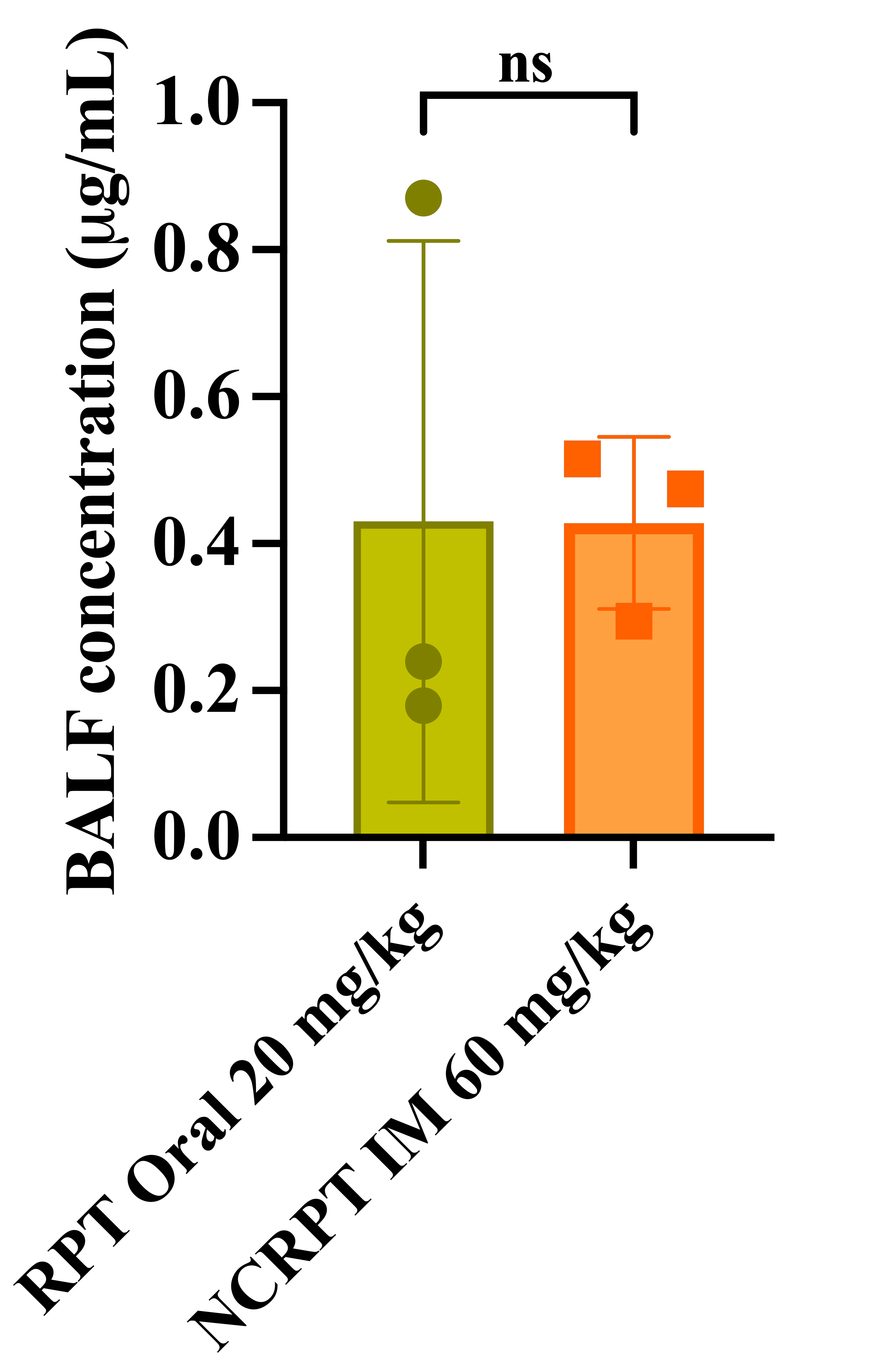

***Figure S6****.* ***BALF pharmacokinetics 24-hour post-dosing****. Data in the graph represent mean ± SD and was analyzed by paired t-test. ns = non-significant (n=3 BALF samples from distinct animals for both groups).*

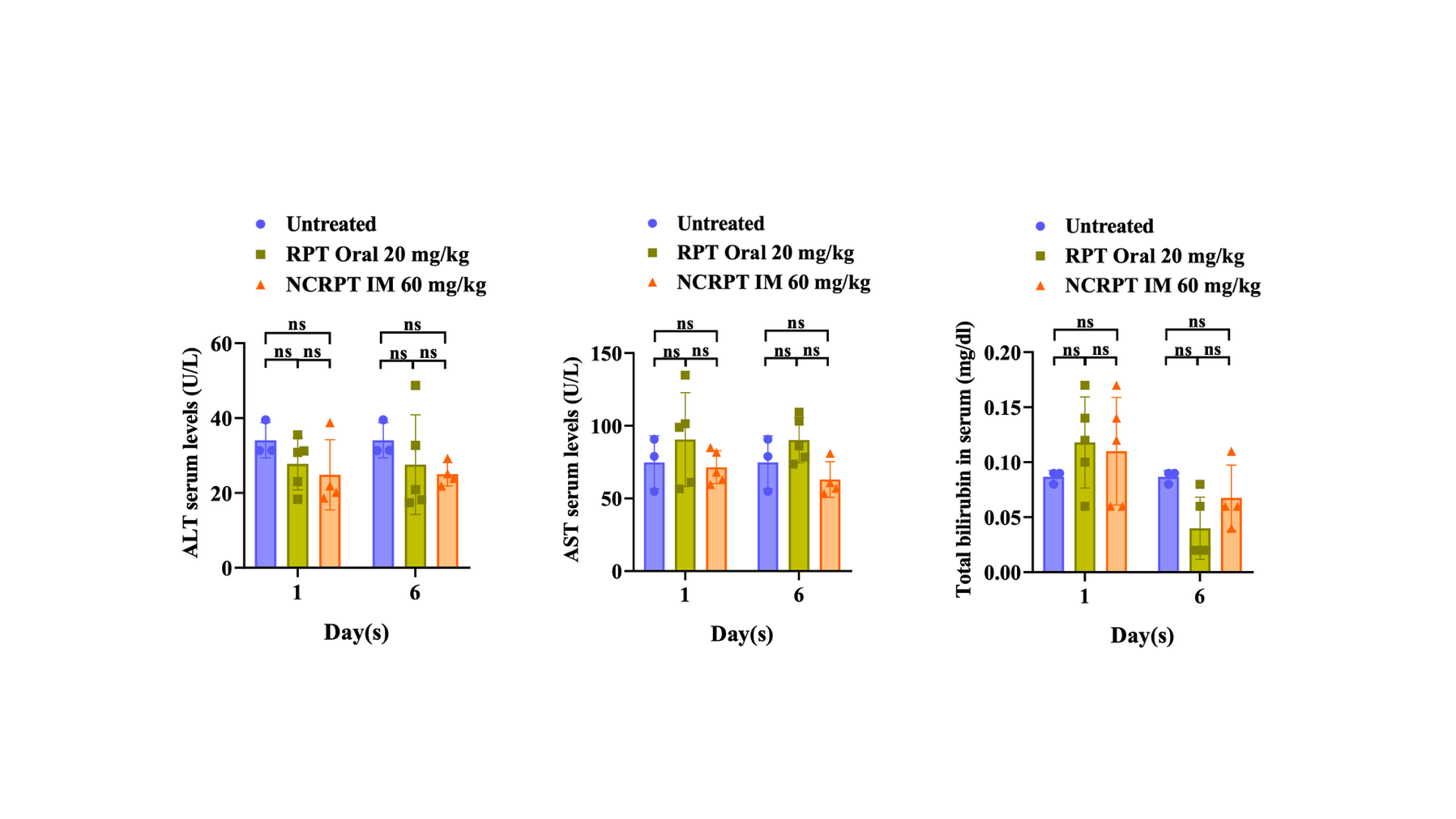

***Figure S7.*** ***NCRPT administration does not cause systemic toxicity****. ALT, AST and total bilirubin levels in serum isolated from untreated, single-dose (20 mg/kg) orally administered RPT and single-dose (60 mg/kg) intramuscularly injected NCRPT on day 1 and day 6 post-drug administration. Data in the graph represent mean ± SD. p-value < 0.05 was considered signiﬁcant. p-values were determined by two-way ANOVA followed by post-hoc Tukey’s multiple comparison tests. No significant difference was observed between any of the groups at either time point for all three tests. (n=3 distinct samples for untreated, n=5 for orally administered, n=5 for NCRPT day 1 and n=4 for NCRPT day 6). One data point from the NCRPT day 1 group for the ALT test was excluded after outlier analysis using the Grubbs test.*

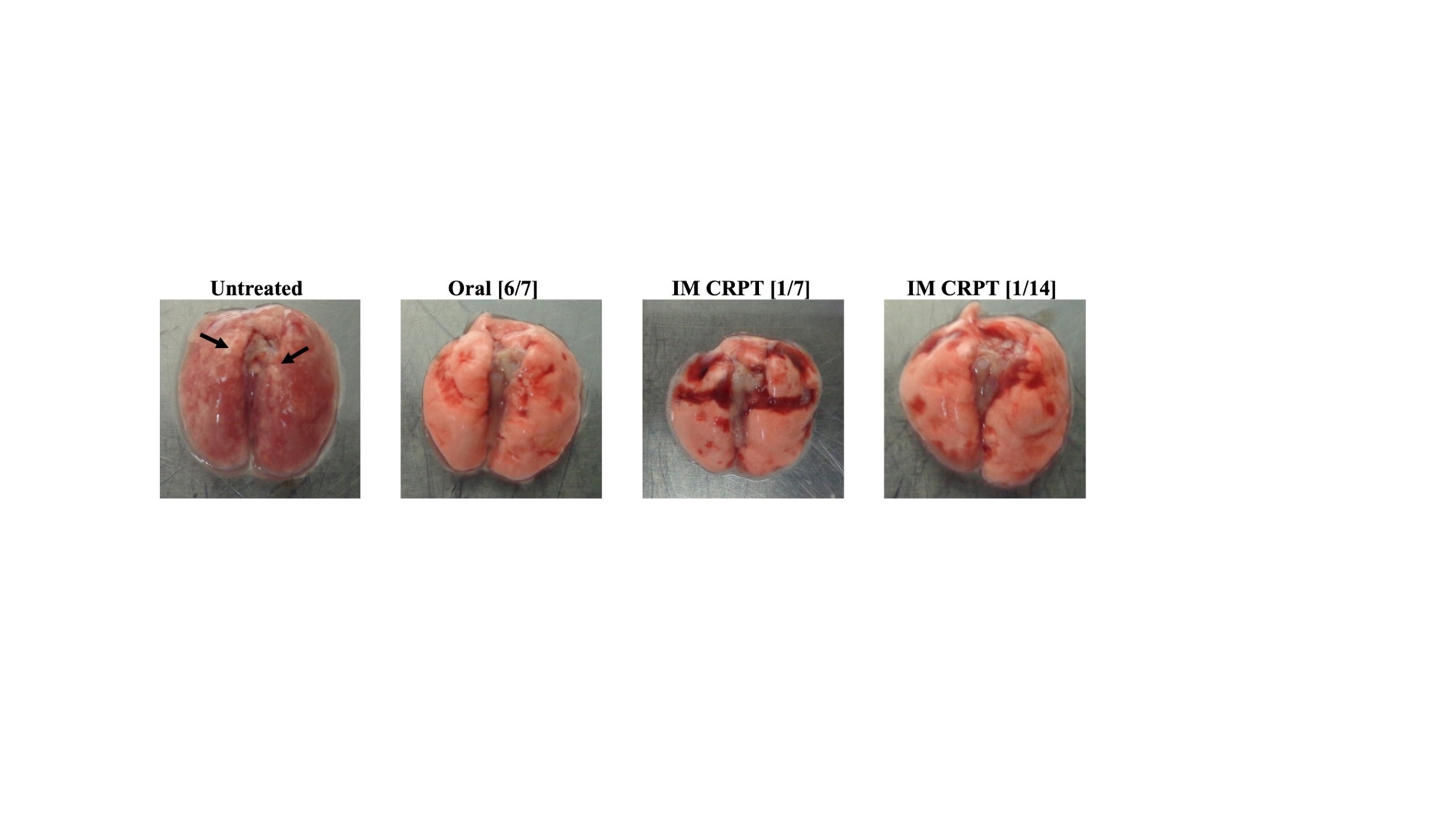

***Figure S8. Lung pathology for pre-exposure prophylaxis efficacy study.*** *Representative images of lungs isolated after four weeks of treatment for pre-exposure prophylaxis efficacy of CRPT in mice infected with Mtb H37Rv. Black arrows indicate representative granulomatous lesions on the lung isolated from the control group mouse.*

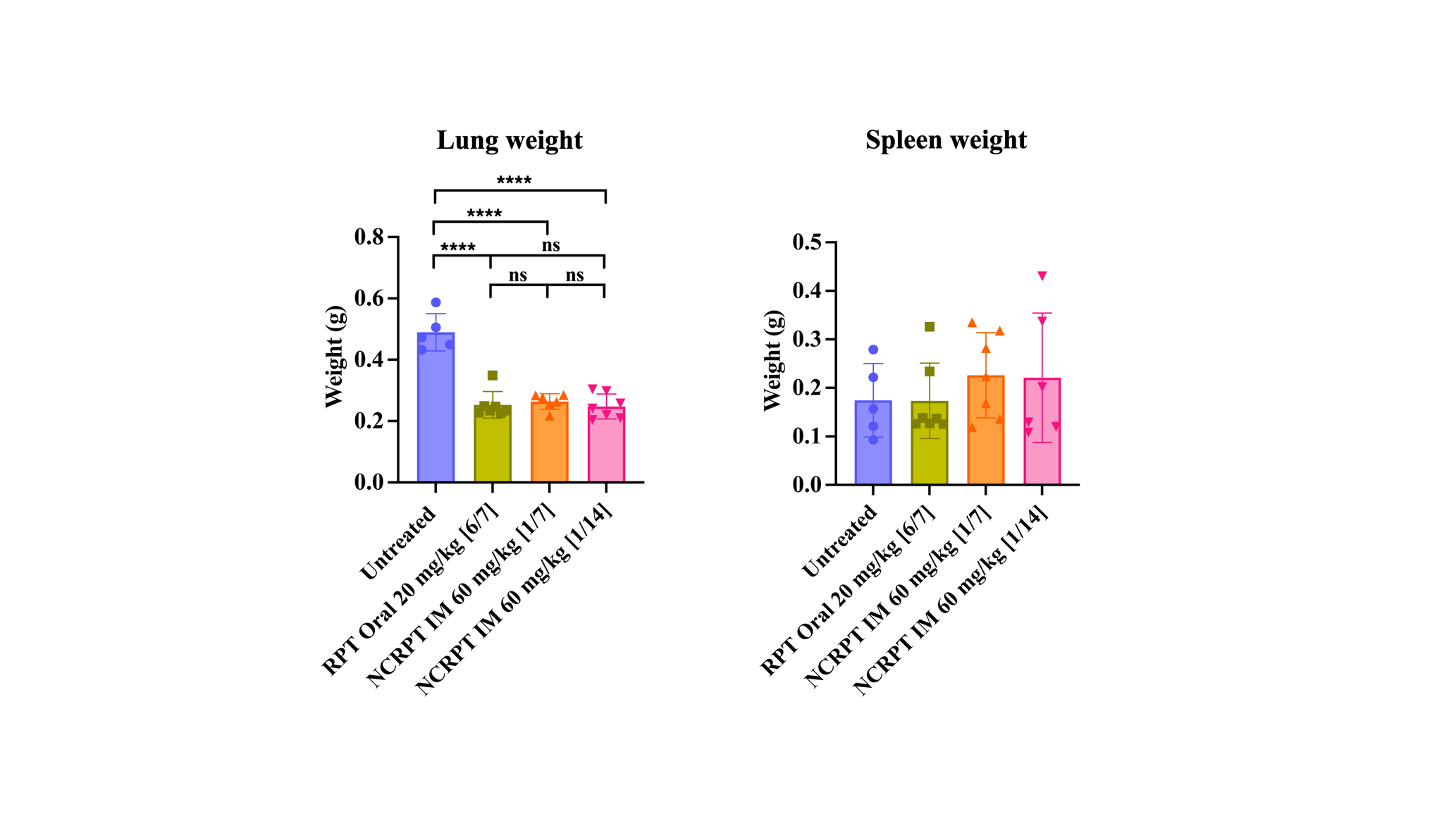

***Figure S9****.* ***Organ weights for pre-exposure prophylaxis efficacy study****. Lung and spleen weights isolated post four weeks of treatment for pre-exposure prophylaxis efficacy of CRPT in mice infected with Mtb H37Rv (n=5 tissues from distinct animals for control, n=7 tissues from distinct animals for all the other groups). Data in the graph represent mean ± SD. p-values were determined by ordinary one-way ANOVA and Tukey’s multiple comparison tests. p-value < 0.05 was considered signiﬁcant. **** p < 0.0001 for control vs oral 20 mg/kg group, ns = non-significant. One data point was excluded for NCRPT 60 mg/kg [1/7] group in lung weight after outlier analysis using Grubbs test.*

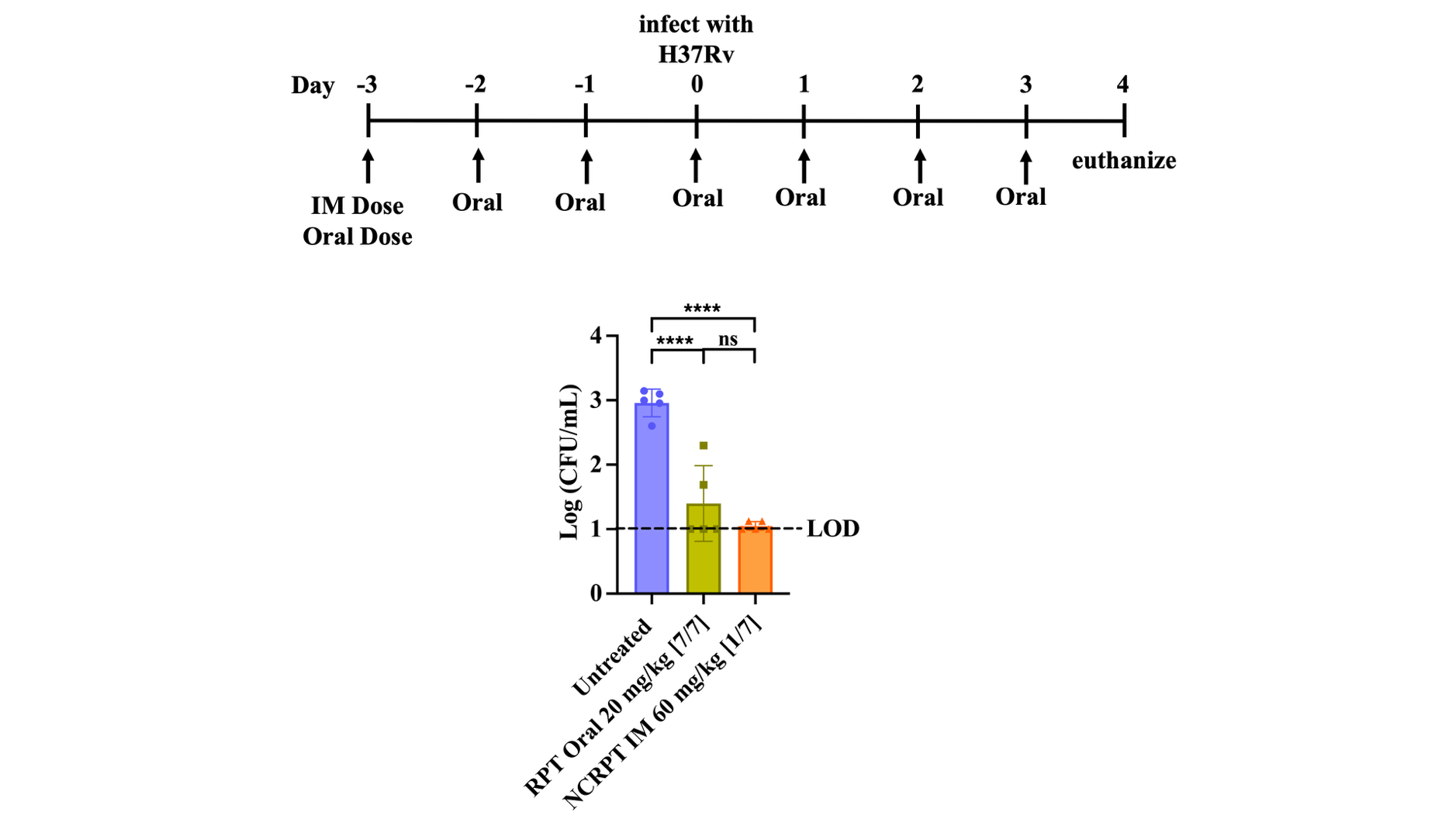

***Figure S10****.* ***Single NCRPT administration has similar efficacy compared to daily oral RPT administration.*** *Bacterial count in the lungs one week after treatment started plotted as Log CFU/mL (n=5 lung samples from distinct animals for all groups). Data in the graph represent mean ± SD. p-values were determined by ordinary one-way ANOVA and Tukey’s multiple comparison tests. p-value < 0.05 was considered signiﬁcant. **** p< 0.0001. ns = non-significant. LOD = limit of detection*

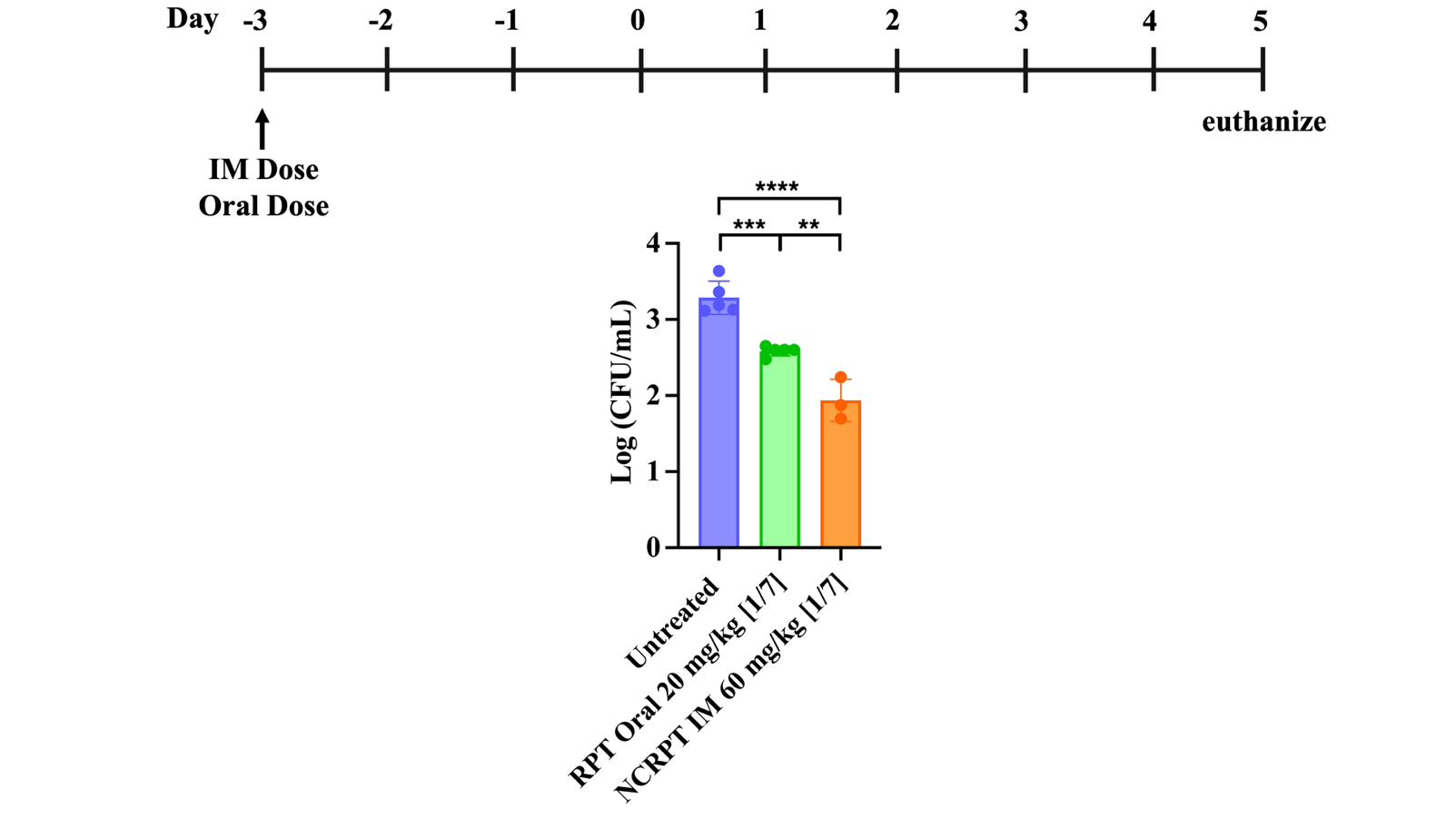

***Figure S11****.* ***Single NCRPT administration has higher efficacy compared to single oral RPT administration.*** *Bacterial count in lungs one week after treatment plotted as Log CFU/mL (n=5 lung samples from distinct animals for control and oral 20 groups and n=3 lung samples from distinct animals for IM CR 60 group). Data in the graph represents mean ± SD. p values were determined by ordinary one-way ANOVA and Tukey’s multiple comparison tests. p-value < 0.05 was considered signiﬁcant. **** p< 0.0001, ***p = 0.0004 for control vs oral 20 group. **p = 0.0023 for oral 20 vs IM CR 60 group.*

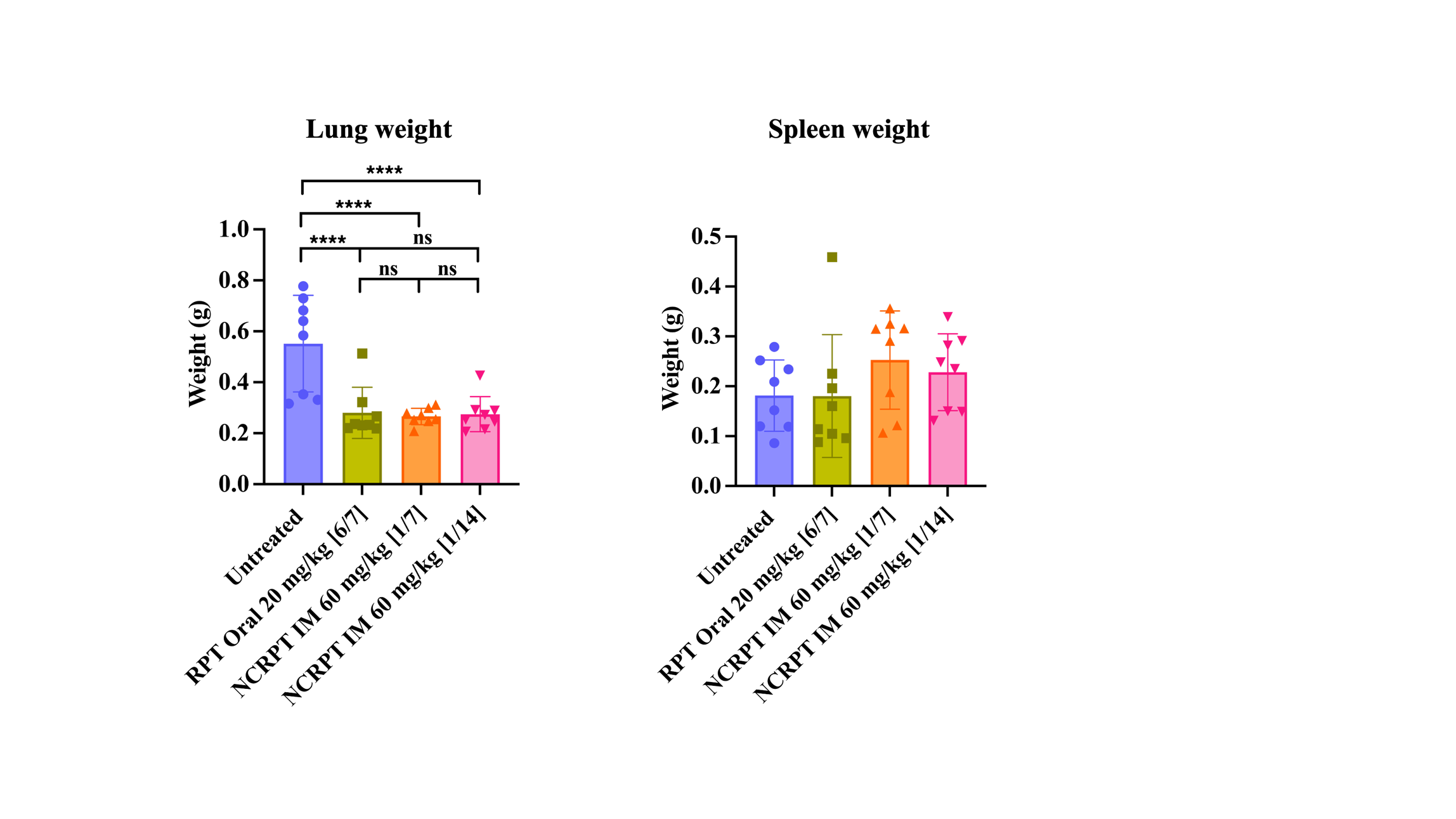

***Figure S12****.* ***Organ weights for therapeutic efficacy study.*** *Lung and spleen weight isolated post four weeks of treatment for the therapeutic efficacy of CRPT in mice infected with Mtb H37Rv (n=8 tissues from distinct animals for all groups). Data in the graph represents mean ± SD. p values were determined by ordinary one-way ANOVA and Tukey’s multiple comparison tests. p-value < 0.05 was considered signiﬁcant. For lung weight, *** p = 0.0003 for control vs oral 20 mg/kg, *** p = 0.0001 for control vs 60 mg/kg [1/7], *** p = 0.0002 for control vs 60 mg/kg [1/14], ns = non-significant.*

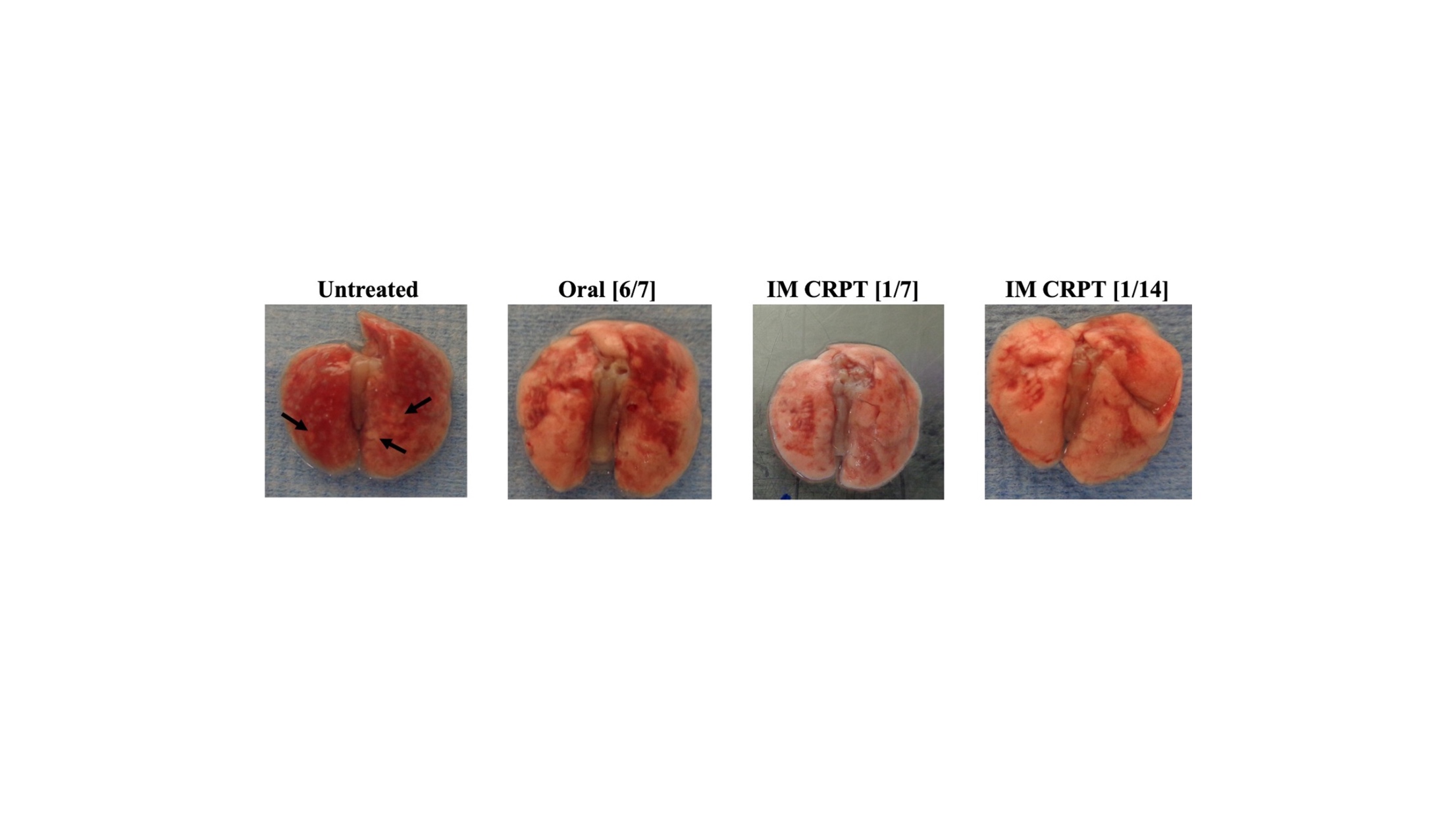

***Figure S13****.* ***Lung pathology for therapeutic efficacy study two weeks post-treatment.*** *Representative images of lungs isolated post two weeks of treatment for the therapeutic efficacy of CRPT in mice infected with Mtb H37Rv. Black arrows indicate representative granulomatous lesions on the lung isolated from the control group mouse.*

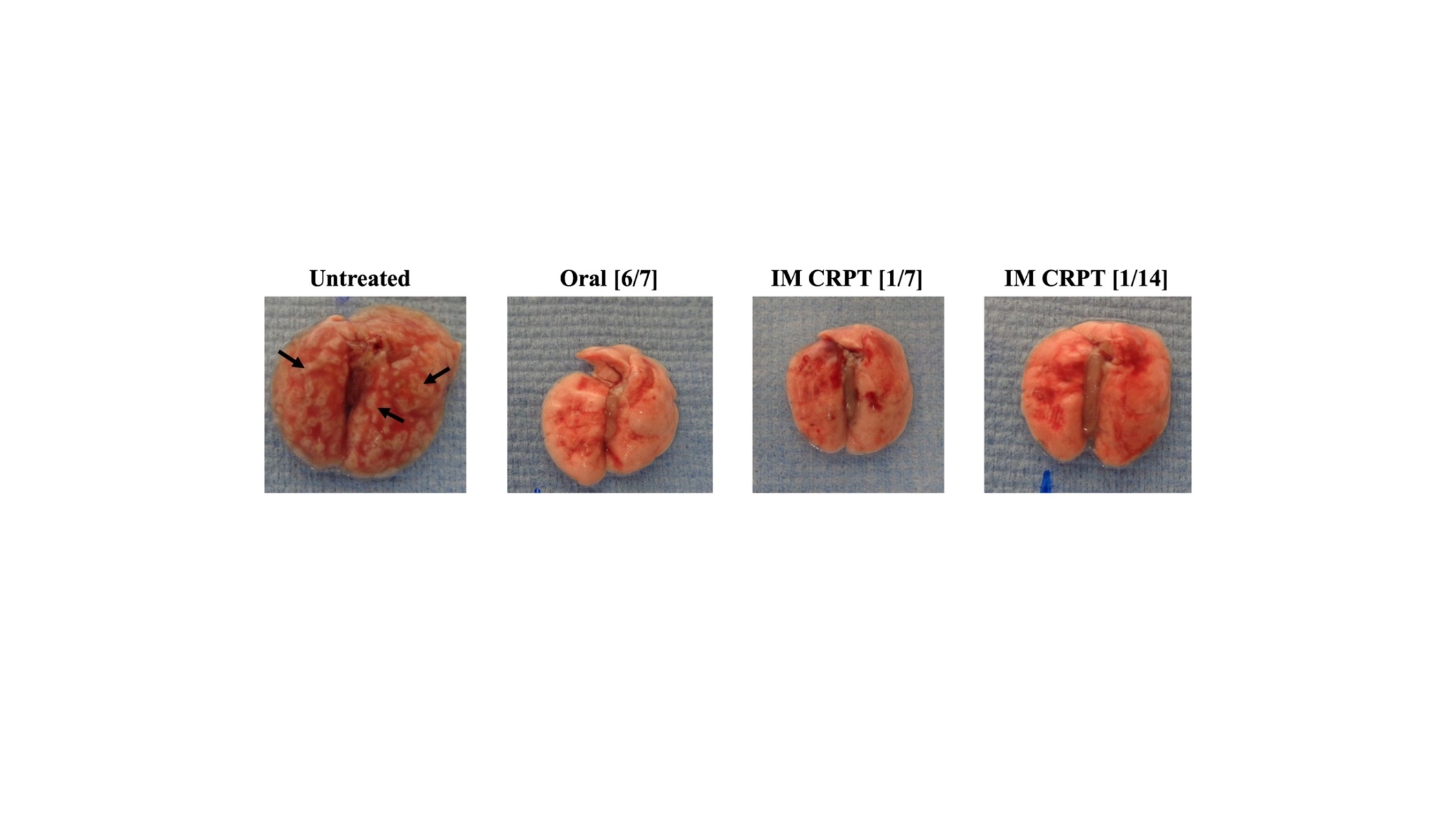

***Figure S14****.* ***Lung pathology for therapeutic efficacy study four weeks post-treatment.*** *Representative images of lungs isolated after four weeks of treatment for the therapeutic efficacy of CRPT in mice infected with Mtb H37Rv. Black arrows indicate representative granulomatous lesions on the lung isolated from the control group mouse.*
